# Bacteriophage susceptibility in clinical *Enterococcus faecalis* reflects host genomic background and Epa-associated phage adsorption

**DOI:** 10.64898/2026.10.12.758625

**Authors:** Tomohiro Nakamura, Jumpei Fujiki, AlcHepNet Investigators, Bernd Schnabl

## Abstract

Cytolysin-positive *Enterococcus faecalis* has been implicated as a disease-associated pathobiont in alcohol-associated hepatitis, motivating development of selective bacteriophage-based strategies. However, strain-level determinants of phage susceptibility remain incompletely understood. We characterized the host range of three *E. faecalis* phages, ΦEf2.1, ΦEf2.2, and ΦEf2.3Ev, across 45 clinical isolates and investigated bacterial and phage factors associated with adsorption and productive infection. Isolates were characterized by whole-genome sequencing, cytolysin status, antimicrobial resistance determinants, and predicted anti-phage defense systems. Phage susceptibility was measured by efficiency of plating (EoP) and adsorption assays. Thirty-five of the 45 isolates (77.8%), including all 15 *cylLs*-positive isolates, were susceptible to at least one of the three phages at EoP ≥0.01. ΦEf2.1 and ΦEf2.2 showed greater activity against *cylLs*-positive isolates, which also carried higher burdens of predicted antimicrobial resistance determinants and anti-phage defense system types. Analysis of 367 complete *E. faecalis* RefSeq genomes classified the variable region of the enterococcal polysaccharide antigen (*epa*) locus into 15 recurrent architecture groups that were associated with phage-specific adsorption patterns. Efficient adsorption did not always result in productive infection, indicating additional post-adsorption barriers. AbiD emerged as a candidate factor associated with reduced ΦEf2.2 productivity, although this signal was prediction-tool dependent, while unexplained post-adsorption blocks suggest the involvement of additional uncharacterized mechanisms. These findings link naturally occurring *epa* architecture to phage adsorption and highlight distinct post-adsorption constraints on host range, providing a framework for rational phage selection against clinically relevant *E. faecalis* populations.

## 1. Introduction

Alcohol-associated hepatitis is a severe inflammatory and cholestatic form of alcohol-associated liver disease and is associated with high short-term mortality (Jophlin et al., 2024). Although alcohol-induced liver injury and host inflammatory responses are central to disease pathogenesis, increasing evidence indicates that the intestinal microbiota contributes to disease progression through the gut-liver axis (Duan et al., 2019). Among microbiota-derived factors, cytolysin-positive *Enterococcus faecalis* has emerged as a clinically relevant pathobiont in alcohol-associated hepatitis (Duan et al., 2019). Cytolysin is a two-subunit exotoxin produced by a subset of *E. faecalis* strains and has hemolytic and bactericidal activity (Van Tyne et al., 2013). The cytolysin locus was originally characterized on the pheromone-responsive plasmid pAD1 and can also be encoded on chromosomal pathogenicity islands or related mobile genetic elements (Gilmore et al., 1994; Shankar et al., 2002; Van Tyne et al., 2013). Duan *et al*. showed that cytolysin-positive *E. faecalis* was increased in patients with alcohol-associated hepatitis and correlated with disease severity and increased mortality (Duan et al., 2019). These findings suggest that cytolysin-positive *E. faecalis* represents a disease-associated bacterial subgroup that may contribute to the pathogenesis and clinical outcome of alcohol-associated hepatitis.

Bacteriophages are viruses that infect bacteria and can selectively kill susceptible bacterial hosts. Their narrow host range can limit broad empirical therapy, but it also helps when the therapeutic goal is to remove a specific pathobiont while preserving the surrounding microbiota (Kortright et al., 2019). This feature is particularly relevant in gut microbiota-associated diseases, where conventional antibiotics may cause broad ecological disruption. Phage-based approaches have therefore been explored as precision tools for editing the intestinal microbiota (Federici et al., 2022; Gan et al., 2023; Hsu et al., 2019). Furthermore, because phage susceptibility is not directly determined by conventional antimicrobial resistance (AMR) genes, phages may provide an alternative or complementary approach for targeting strains carrying AMR determinants (Dedrick et al., 2019; Pirnay et al., 2024; Schooley et al., 2017). In the context of alcohol-associated liver disease, bacteriophages targeting cytolytic *E. faecalis* reduced cytolysin burden and attenuated ethanol-induced liver disease in microbiota-humanized mouse models (Duan et al., 2019). In addition, ΦEf2.1 and ΦEf2.2 have been used as candidate phages targeting cytolytic *E. faecalis*, and our recent work showed that resistance to these phages can be accompanied by fitness costs, including impaired intestinal colonization (Fujiki et al., 2025). In addition, ΦEf2.3Ev, an evolved derivative of ΦEf2.3 isolated following co-culture with *E. faecalis* strain efSD10, may represent another promising candidate for targeting cytolytic *E. faecalis* (Fujiki et al., 2026). ΦEf2.1, ΦEf2.2, and ΦEf2.3Ev have been selected for evaluation as an oral phage therapy targeting *cylLs*-positive *E. faecalis* in patients with alcohol-associated hepatitis (ClinicalTrials.gov identifier: NCT06750588). However, the lytic activity and host range of ΦEf2.1, ΦEf2.2, and ΦEf2.3Ev across diverse clinical *E. faecalis* isolates have not yet been systematically evaluated.

A major determinant of phage host range is the interaction between the phage and bacterial surface receptors. In *E. faecalis*, the enterococcal polysaccharide antigen, Epa, has emerged as a key receptor-associated cell surface polymer for multiple *E. faecalis* phages (Al-Zubidi et al., 2019; Chatterjee et al., 2020; Fujiki et al., 2025; Ho et al., 2018). Epa is a complex cell wall polysaccharide that contributes to cell envelope integrity, adhesion, bile salt resistance, intestinal colonization, and virulence-related phenotypes (Guerardel et al., 2020; Rigottier-Gois et al., 2015). The *epa* locus contains conserved genes involved in synthesis of the core rhamnopolysaccharide backbone, as well as a variable region that encodes strain-specific surface decorations (Alrafaie et al., 2024; Guerardel et al., 2020). Previous studies have shown that mutations in *epa* genes can reduce phage adsorption or confer phage resistance (Al-Zubidi et al., 2019; Chatterjee et al., 2020; Ho et al., 2018). Importantly, recent work showed that the *epa* variable region itself can be required for *Enterococcus* phage infection: deletion of the V583 *epa* variable locus abolished infection by phage SHEF13, and complementation with the V583 *epa* variable region restored infection (Alrafaie et al., 2024). In cytolytic *E. faecalis* EF01-derived phage-resistant variants, defects affecting Epa biosynthesis, including an *epaR* mutation and loss of *galE*, reduced adsorption of ΦEf2.1 and ΦEf2.2, suggesting that Epa-associated surface structures are important for recognition by these phages (Fujiki et al., 2025). However, the broader diversity of the *epa* variable region among *E. faecalis* genomes, and whether naturally occurring *epa* variable-region patterns predict adsorption specificity of phages targeting clinical isolates, remains poorly understood.

Phage adsorption alone may not be sufficient to support productive infection. After attachment to the bacterial surface, infection can still be blocked at post-adsorption steps such as genome entry, replication, transcription, or assembly. Bacteria encode diverse anti-phage defense systems, including restriction-modification systems, abortive infection systems, CRISPR-Cas systems, retrons, and other defense modules (Bernheim and Sorek, 2020; Payne et al., 2022; Tesson et al., 2022). Recent studies in enterococci have highlighted the importance of type IV restriction enzymes in phage defense. For example, a type IV restriction enzyme encoded on an *E. faecalis* mobilizable plasmid, TIV-RE, restricts phage replication, while an enterococcal phage evolved a TifA anti-restriction protein that counteracts this defense (Bullen et al., 2024). Despite these advances, the contribution of predicted defense systems to phage productivity across *E. faecalis* clinical isolates remains incompletely understood.

In this study, we evaluated how effectively three *E. faecalis* phages, ΦEf2.1, ΦEf2.2, and the evolved phage ΦEf2.3Ev derived from ΦEf2.3, target clinical *E. faecalis* isolates, with particular emphasis on cytolysin-positive isolates derived predominantly from patients with alcohol-associated hepatitis. We first characterized cytolysin status, hemolytic activity, phylogenetic relationships, and antimicrobial resistance gene content among the isolates. We then measured phage lytic activity and adsorption efficiency across this strain collection. To determine whether receptor-associated variation contributes to phage activity, we classified *epa* variable regions from complete *E. faecalis* RefSeq genomes into super-patterns and assigned clinical isolates to these groups. Finally, we explored whether predicted anti-phage defense systems were associated with reduced phage productivity among isolates that retained efficient adsorption.

## 2. Materials and Methods

### 2.1. Fecal samples

Fecal samples from healthy controls, heavy drinkers and patients with alcohol-associated hepatitis were collected through the AlcHepNet between April 2019 and November 2023 at 9 clinical centers in the United States: Cleveland Clinic, Indiana University, Mayo Clinic, University of Louisville, Beth Israel Deaconess Medical Center and University of Massachusetts, University of Pittsburgh, University of Texas Southwestern, and Virginia Commonwealth University. Inclusion criteria included a clinical diagnosis of alcohol-associated hepatitis, serum total bilirubin >3 mg/dL, age ≥21 years, and the ability to provide written informed consent. Re-enrollment of alcohol-associated hepatitis patients was permissible if a new episode of alcohol-associated hepatitis occurred at least 24 weeks after the most recent enrollment; however, no re-enrolled cases were included in this study. Exclusion criteria included hemochromatosis, autoimmune liver disease, Wilson disease, non-alcoholic fatty liver disease, viral hepatitis, pregnancy, or inability to comply with study procedures.

### 2.2. Bacterial strains and bacteriophages

The laboratory *Enterococcus faecalis* strain OG1RF (ATCC 47077) was obtained from the American Type Culture Collection (ATCC, Manassas, VA, USA). *E. faecalis* V583, a vancomycin-resistant clinical isolate, was kindly provided by Dr. Michael S. Gilmore (Shankar et al., 2002). The cytolytic *E. faecalis* clinical isolate EF01 was previously isolated from a fecal sample obtained from a patient with alcohol-associated hepatitis and characterized in our previous study (Duan et al., 2019). The additional clinical *E. faecalis* isolates analyzed in the present study were newly isolated from fecal samples collected through the AlcHepNet cohort, including patients with alcohol-associated hepatitis, heavy drinkers, and healthy controls. (AlcHepNet). The *E. faecalis* bacteriophages ΦEf2.1, ΦEf2.2, and ΦEf2.3 were previously isolated from sewage and characterized in our previous study (Duan et al., 2019). ΦEf2.3Ev is an evolved derivative of ΦEf2.3 that was isolated following co-culture of ΦEf2.3 with *E. faecalis* strain efSD10 (Fujiki et al., 2026). According to the current viral taxonomy accessed in August 2026, ΦEf2.1 and ΦEf2.3 are classified within the genus *Kochikohdavirus*, whereas ΦEf2.2 is classified within the genus *Saphexavirus*. For preparation of high-titer phage stocks, EF01 was used as the propagation host. Phages were amplified using either the double-layer agar (DLA) method or liquid-culture amplification, as appropriate.

### 2.3. Isolation and qPCR screening of cytolysin-positive Enterococcus faecalis

Fecal samples from 194 participants enrolled in the AlcHepNet, including patients with alcohol-associated hepatitis, heavy drinkers, and healthy controls, were screened for *E. faecalis* isolation. Detection of cytolysin-positive *Enterococcus faecalis* colonies was performed as previously described, with modifications (Cabré et al., 2023). Briefly, approximately 10 mg of fecal sample was suspended in 3 mL of Enterococcosel broth (Becton Dickinson, Franklin Lakes, NJ, USA) and streaked onto Enterococcosel agar plates and incubated at 37°C overnight. Multiple colonies were picked and transferred onto Tryptic Soy Agar (Becton Dickinson) with 5 % horse blood (Thermo Fisher Scientific, Waltham, MA, USA) plates to generate master plates. Colony quantitative polymerase chain reaction was performed to detect an *E. faecalis*-specific 16S rRNA gene target and the cytolysin-associated gene *cylLs* using the following primer pairs: *E. faecalis* 16S rRNA gene, forward 5′-CGCTTCTTTCCTCCCGAGT-3′ and reverse 5′-GCCATGCGGCATAAACTG-3′ (expected amplicon size, 142 bp); and *cylLs*, forward 5′-GTAAAATAAGTAAAATCAAGAAAACTATTACTC-3′ and reverse 5′-CAAAAGAAGGACCAACAAGTTCTAATT-3′ (expected amplicon size, 61 bp). Reactions were performed using Fast SYBR^TM^ Green Master Mix (Thermo Fisher Scientific) on an Applied Biosystems^TM^ StepOnePlus^TM^ real-time PCR system (Thermo Fisher Scientific). Each reaction contained 10 μL of Fast SYBR^TM^ Green Master Mix, 1 μL of primer mix (forward and reverse, 10 μM each), and 9 μL of nuclease-free water. A single colony picked from the horse blood agar master plate was then suspended directly into the reaction mixture and used as the template. The cycling conditions were 95 °C for 3 min for initial denaturation, followed by 40 cycles of 95 °C for 15 s for denaturation and 60 °C for 1 min for annealing and extension. Colonies were classified as *E. faecalis* positive or cytolysin positive when the *E. faecalis* 16S rRNA gene or *cylLs* target was detected with Ct values of ≤30 or ≤32, respectively, and showed a single melting-curve peak at the expected melting temperature. Selected single colonies were inoculated into Brain Heart Infusion (BHI) broth (Becton Dickinson) and incubated at 37°C for 18 h. Cultures were mixed with sterile glycerol to a final concentration of 25 % and stored at −80 °C.

### 2.4. Assessment of hemolytic activity on horse blood agar

Beta-hemolytic activity was evaluated on horse blood agar plates as a phenotypic readout of cytolysin activity (Reynolds et al., 2025). Overnight cultured bacterial strains in BHI broth were streaked onto TSA (Becton Dickinson) with 5 % horse blood agar (Thermo Fisher Scientific) and incubated at 37 °C for 18 hours. Beta-hemolysis was defined as a clear zone surrounding bacterial growth and was scored as present or absent by visual inspection.

### 2.5. Whole-genome sequencing, assembly, and annotation

Genomic DNA from all clinical isolates was extracted using the DNeasy PowerLyser PowerSoil Kit (QIAGEN, Venlo, Netherlands). The extracted DNA was then subjected to short-read sequencing on the Illumina NovaSeq X Plus platform or NextSeq 2000 platform with 2 × 150 bp paired-end reads. Reads were quality filtered and adaptor clipped using Trimmomatic v0.39 (Bolger et al., 2014) (LEADING:20 TRAILING:20 SLIDINGWINDOW:4:15, MINLEN 80) and assembled de novo using Shovill v1.1.0 with default settings (Seemann T (2016) *Shovill* Github https://github.com/tseemann/shovill). Genome assemblies were annotated with Bakta v1.12.0 (database: v6.0, full) (Schwengers et al., 2021).

### 2.6. Basic bioinformatics

Multi-locus sequence types (MLST) were assigned using mlst v 2.32.2 (Seemann T, *mlst,* Github https://github.com/tseemann/mlst) (Jolley et al., 2018) with the *E. faecalis* MLST scheme. Isolates for which a sequence type could not be assigned were labeled as NA in the figures. The Cytolysin operon was identified in assembled genomes using ABRicate v1.2.0 (Seemann T, *Abricate*, Github https://github.com/tseemann/abricate) against the VFDB database (2025-Dec-5) (Chen et al., 2016). AMR determinants were identified from genome assemblies using StarAMR v0.12.1 (Bharat et al., 2022). The AMR determinant count was defined as the number of unique AMR genes and resistance-associated mutations reported by StarAMR. Anti-phage defense systems were predicted by DefenseFinder v2.0.0 (Tesson et al., 2022) and PADLOC v2.0.0 (Payne et al., 2021). To assess the presence of the recently described *E. faecalis* type IV restriction enzyme TIV-RE, the amino acid sequence of EF_B0059 was used as the query for a targeted homology search (Bullen et al., 2024). Predicted protein sequences from all clinical isolate genomes, annotated using Bakta, were combined into a single protein database and searched using BLASTP v2.17.0+ (Camacho et al., 2009). Hits were considered candidate EF_B0059 homologs when they met all of the following thresholds: amino acid identity ≥35%, query coverage ≥70%, and E-value ≤1 × 10⁻⁵. For each isolate, the best-scoring hit passing these criteria was retained, and the presence or absence of a TIV-RE homolog was summarized at the isolate level.

### 2.7. Comparative analysis of tail-associated genomic regions of ΦEf2.1 and ΦEf2.3Ev

The genome sequences of ΦEf2.1 (MK693030) and ΦEf2.3Ev were used for comparative analysis. The ΦEf2.3Ev genome sequence was reconstructed *in silico* from the parental ΦEf2.3 reference genome (MK721192) by incorporating the experimentally identified ΦEf2.3Ev mutation. Both phage genomes were re-annotated using Pharokka v1.8.1 (Bouras et al., 2023), followed by structure-informed functional annotation with Phold v1.1.0 (Bouras et al., 2026). Based on the resulting functional annotations, the genomic regions containing the tail-associated gene clusters were identified. For each phage, the comparison region was defined as the contiguous genomic interval bounded by coding sequences (CDSs) annotated with tail-associated functions at both ends. The corresponding regions from ΦEf2.1 and ΦEf2.3Ev were then compared and visualized using LoVis4u v0.1.6 (Egorov and Atkinson, 2025) to assess conservation of gene organization.

### 2.8. Core-genome phylogeny and visualization

A core-genome alignment was generated using Panaroo v1.6.0 (Tonkin-Hill et al., 2020) with the following options: --clean-mode strict, --remove-invalid-genes, -a core, --aligner mafft (Katoh and Standley, 2013), and --core_threshold 0.95. The resulting filtered core-gene alignment was used for phylogenetic inference. A maximum-likelihood phylogeny was inferred using IQ-TREE version 3.0.1 (Wong et al., 2026). The best-fit nucleotide substitution model was selected using ModelFinder (Kalyaanamoorthy et al., 2017), and branch support was assessed using 1,000 SH-aLRT replicates (Guindon et al., 2010) and 1,000 ultrafast bootstrap replicates(Hoang et al., 2018). The final phylogeny used for visualization included the 45 clinical isolates and three additional reference or laboratory strains, EF01, OG1RF, and V583. The tree was visualized and annotated using iTOL v7 (Letunic and Bork, 2024).

### 2.9. Construction of the RefSeq epa super-pattern (SP) catalog and representative selection

All *E. faecalis* genome FASTA files registered as complete genomes in NCBI as of April 9, 2026—a total of 376 files—were downloaded and re-annotated using Bakta v1.12.0 (database: v6.0, full). For each annotated GenBank file, predicted proteins were extracted with Biopython v1.86 (Cock et al., 2009) and searched against a V583-derived *epa* query set using BLASTP v2.17.0+. The query set contained proteins encoded by the V583 *epa* core region (EF2177– EF2198) and the *epa* variable region (EF2164–EF2176), and the EF2163 homolog as a terminal boundary anchor. BLASTP hits were retained using an e-value cutoff of ≤1e-5, minimum amino-acid identity of 25%, and minimum query coverage of 50%. For downstream architecture-based SP classification, the *epaR* (EF2177)-to-EF2163 intervals from the 367 RefSeq genomes that met complete-locus quality control criteria were used; loci exhibiting missing EF2163 anchors, strand inconsistencies, incomplete boundary recovery, or >20 CDSs between *epaR* and the EF2163 homolog were excluded. CDS protein sequences were clustered into orthogroups using MMseqs2 v18.8cc5c (Steinegger and Söding, 2017) easy-cluster with the following parameters: --min-seq-id 0.50, --c 0.80, and --cov-mode 1. Each locus was converted into an ordered orthogroup string, and identical strings were collapsed into exact patterns. To capture broader architectural similarity beyond exact identity, exact patterns were further grouped into super-patterns (SPs) using a weighted architecture distance based on orthogroup-content Jaccard distance and orthogroup-adjacency Jaccard distance, with weights of 0.70 and 0.30, respectively. Exact patterns connected at a distance of ≤0.30 were assigned to the same SP. The resulting set of SPs constituted the RefSeq *epa* SP catalog. For each SP, the representative exact pattern was selected by the lowest mean within-SP architecture distance, and the representative sample was then selected from that exact pattern by CDS count closest to the exact-pattern median. In the event of a tie, the smaller CDS count was used to break the tie, and the lexicographic accession order was used to resolve any additional ties.

### 2.10. Architecture-distance tree of RefSeq epa variable-region loci

An architecture-distance tree was constructed from the RefSeq SP catalog ordered orthogroup profile table, which contained the sample identifier, CDS count, exact-pattern assignment, SP assignment, and ordered orthogroup string for each QC-passing RefSeq *epaR*-to-EF2163 locus. Pairwise architecture distances were calculated from the ordered orthogroup strings as the weighted sum of orthogroup-content Jaccard distance and orthogroup-adjacency Jaccard distance, using weights of 0.70 and 0.30, respectively. A neighbor-joining tree was constructed from the resulting distance matrix using the DistanceTreeConstructor module in Biopython v1.86 and exported in Newick format. The tree was visualized in iTOL v7.

### 2.11. Assignment of clinical-strain epa loci to RefSeq SPs

Clinical-strain *epaR*-to-EF2163 loci generated using the same extraction and QC procedure described above were assigned to the RefSeq SP catalog. Complete clinical-strain loci were converted to ordered orthogroup strings and assigned to RefSeq exact patterns and SPs using the same orthogroup framework and architecture-distance metric used for RefSeq SP catalog construction; query CDSs were assigned to RefSeq orthogroups by BLASTP with e-value ≤1e-5, amino-acid identity ≥35%, and query coverage ≥70%. Clinical strains without complete *epaR*-to-EF2163 recovery were analyzed using recovered partial or terminal *epa* fragments. The EF2163-containing contig was identified from filtered BLASTP hits generated during initial extraction, and the EF2163 homolog was used as a terminal anchor to extract an EF2163-proximal GenBank fragment from the selected terminal-fragment orientation. Subsequently, partial or terminal fragments were compared with all RefSeq SP representative loci by BLASTP. CDS protein sequences were extracted from the query and representative GenBank files, and a separate protein BLAST database was constructed for each representative locus. Query CDS protein sequences were searched against each representative database using BLASTP with an e-value cutoff of 1e-5, amino-acid identity ≥25%, and query coverage ≥50%. For each query-representative pair, similarity was summarized as the number of matched query CDSs, the fraction of query CDSs matched, the total bitscore, the normalized bitscore per query CDS, the mean bitscore per matched query CDS, the mean percent identity, and the mean query coverage. The top-ranked representative SP was assigned when supported by locus-architecture visualization.

### 2.12. Visualization of epa locus architectures

Representative and clinical-strain *epa* variable-region locus architectures were visualized using pyGenomeViz v1.6.1 (Shimoyama Y, *pyGenomeViz*, Github https://github.com/moshi4/pyGenomeViz). CDS colors and labels were assigned from the curated global orthogroup annotation table, and links between loci were drawn only for CDSs assigned to the same curated orthogroup. For the representative SP architecture figure, one RefSeq representative locus was shown for each SP001–SP015 and ordered by SP number. For SP-specific clinical-strain visualizations, the RefSeq representative was shown first, followed by complete assignments and then partial or terminal assignments.

### 2.13. Efficiency of plating assay and phage adsorption assay

The lytic activity and host range of ΦEf2.1, ΦEf2.2, and ΦEf2.3Ev against *E. faecalis* strains were evaluated by efficiency of plating (EoP) assays with modifications from a previously described method (Fujiki et al., 2022). Each *E. faecalis* strain was cultured overnight in BHI broth at 37°C. The bacterial culture was mixed with BHI top agar containing 0.5% agarose (Thermo Fisher Scientific) and overlaid onto a BHI agar plate to generate a bacterial lawn. Serially, 2.5 μL of diluted phage suspensions were then spotted onto the bacterial lawns, and the plates were incubated overnight at 37°C. Plaques were counted at an appropriate dilution, and the phage titer was expressed as plaque-forming units per milliliter (PFU/mL). EoP was calculated as the phage titer obtained on each test strain divided by the phage titer obtained on EF01, which was used as the reference host for all three phages. Each assay was performed in triplicate, and the mean EoP value from the three replicates was used for downstream analyses. For subsequent analyses, isolates with EoP ≥0.1 were classified as sensitive, those with 0.01 ≤ EoP <0.1 as intermediate, and those with EoP <0.01 as resistant. Phage adsorption was assessed with modifications from a previously described method (Fujiki et al., 2021). Each *E. faecalis* strain was cultured overnight in 5 mL of BHI broth at 37°C. 50 μL of the overnight bacterial culture was mixed with 50 μL of phage suspension at a multiplicity of infection (MOI) of approximately 0.1. The mixture was incubated at room temperature for 10 min to allow phage adsorption, then immediately centrifuged at 10,000 × g for 10 min at 4°C. The resulting supernatant, containing unadsorbed phage particles, was collected and serially diluted in SM buffer (10 mM MgSO_4_, 100 mM NaCl, 0.01% gelatin, and 50 mM Tris-HCl, pH 7.5). 2.5 μL of each dilution was spotted onto BHI agar plates overlaid with BHI top agar containing EF01, and the plates were incubated overnight at 37°C. The titer of unadsorbed phage in the supernatant was determined by plaque counting. The adsorption rate was calculated as: Adsorption (%) = [1 − (titer of unadsorbed phage / initial phage titer)] × 100. Each adsorption assay was performed in triplicate, and the mean adsorption rate from the three replicates was used for downstream analyses. For subsequent analyses, adsorption rates were categorized as low (<30%), intermediate (30% to <80%), or high (≥80%).

### 2.14. Statistical analysis

Statistical analyses were performed using R v4.5.3 (2026-03-11). AMR determinant counts were compared between *cylLs*-positive (n = 15) and *cylLs*-negative (n = 30) clinical isolates using a Wilcoxon rank-sum test. EoP values and adsorption efficiencies were compared between *cylLs*-positive and *cylLs*-negative clinical isolates separately for each phage using Wilcoxon rank-sum tests. For each outcome, *P* values from the three phage-specific comparisons were adjusted using the Benjamini–Hochberg (BH) method. For analyses stratified by *epa* SP, comparisons were restricted to SPs containing at least three strains. Differences in adsorption efficiency and EoP among the three phages were evaluated separately using Friedman tests, with measurements from the same strain treated as matched observations. Pairwise comparisons between phages were performed using paired Wilcoxon signed-rank tests, and the three pairwise *P* values were adjusted using the BH method separately within each SP and outcome. The numbers of distinct predicted anti-phage defense system types were compared between *cylLs*-positive (n = 17) and *cylLs*-negative (n = 31) strains across the full 48-strain collection using Wilcoxon rank-sum tests, separately for DefenseFinder and PADLOC. Associations between individual predicted defense systems and infection outcome were assessed using Fisher’s exact tests. For each phage and prediction tool, these analyses were restricted to strains with adsorption efficiencies ≥80%, and defense system presence or absence was compared between strains supporting productive infection (EoP ≥0.1) and those showing strong post-adsorption blocking (EoP <0.01). Strains with intermediate EoP values (0.01 ≤ EoP <0.1) were excluded from these comparisons. *P* values were adjusted across the evaluated defense systems using the BH method separately for each phage and prediction tool. EoP values were compared between AbiD-positive and AbiD-negative strains using Wilcoxon rank-sum tests separately for each phage and prediction tool. Wilcoxon rank-sum tests, Wilcoxon signed-rank tests, and Fisher’s exact tests were two-sided. *P* values <0.05 were considered statistically significant, with BH-adjusted *P* values used where adjustment for multiple testing was specified.

## 3. Results

### 3.1. Cytolysin-positive E. faecalis clinical isolates carry a higher burden of predicted antimicrobial resistance determinants

We first isolated *E. faecalis* from all 194 fecal samples available from the AlcHepNet cohort. This cohort included samples from patients with alcohol-associated hepatitis, heavy drinkers, and healthy controls. Candidate colonies were recovered using a selective medium and subsequently screened by quantitative polymerase chain reaction to confirm the presence of *E. faecalis* and to detect the cytolysin-associated gene *cylLs*. We isolated 45 *E. faecalis* strains, of which 15 carried *cylL*s (Table S1). Subsequently, these isolates were subjected to genomic and phenotypic characterization with the clinical isolate EF01, a host for phage amplification, and the laboratory *E. faecalis* strains OG1RF and V583. The qPCR-based *cylLs* detection results exhibited substantial concordance with the NGS-based detection of the cytolysin operon and with hemolytic activity on blood agar (Fig. 1A and Fig. S1A). The clinical isolate collection included diverse sequence types, but *cylLs*-positive isolates were concentrated in a limited number of multilocus sequence types (STs) (Fig. 1A and 1B). In particular, *cylLs*-positive isolates were enriched in ST179 (7/7), ST16 (4/4), and ST6 (2/3). Furthermore, the phylogenetic tree showed that ST179 and ST16 are closely related. We next compared antimicrobial resistance determinant burden between *cylLs*-positive and *cylLs*-negative isolates. *cylLs*-positive isolates carried a significantly higher number of antimicrobial resistance determinants than *cylLs*-negative isolates (median, 5 versus 1 AMR determinants) (Fig. 1C). Consistent with this higher AMR determinant burden, several resistance determinants were enriched among *cylLs*-positive isolates (Fig. 1A and Fig. S1B). Genotype-based predictions of antimicrobial resistance phenotypes are summarized in Fig. S1C. The tetracycline resistance gene *tet(M)* was detected in all *cylLs*-positive isolates, whereas it was present in only a minority of *cylLs*-negative isolates. The macrolide resistance gene *erm(B)* was also frequent in the *cylLs*-positive group, as were aminoglycoside resistance determinants, including *aac(6’)-aph(2’’)*, *ant(6)-Ia*, and *aph(3’)-III*. In addition, all clinical isolates classified as ST6 harbored mutations in the quinolone resistance-determining regions *gyrA* (S83I) and *parC* (S80I). Among them, the EF9c1 strain was the only one out of 45 to possess the VanHAX cassette, which is associated with vancomycin resistance. In contrast, *lsa(A)* was detected broadly across both *cylLs*-positive and *cylLs*-negative isolates. Together, these data show that *cylLs*-positive *E. faecalis* isolates in this collection are concentrated in specific sequence types and carry a higher burden of antimicrobial resistance determinants.

**Figure 1.**
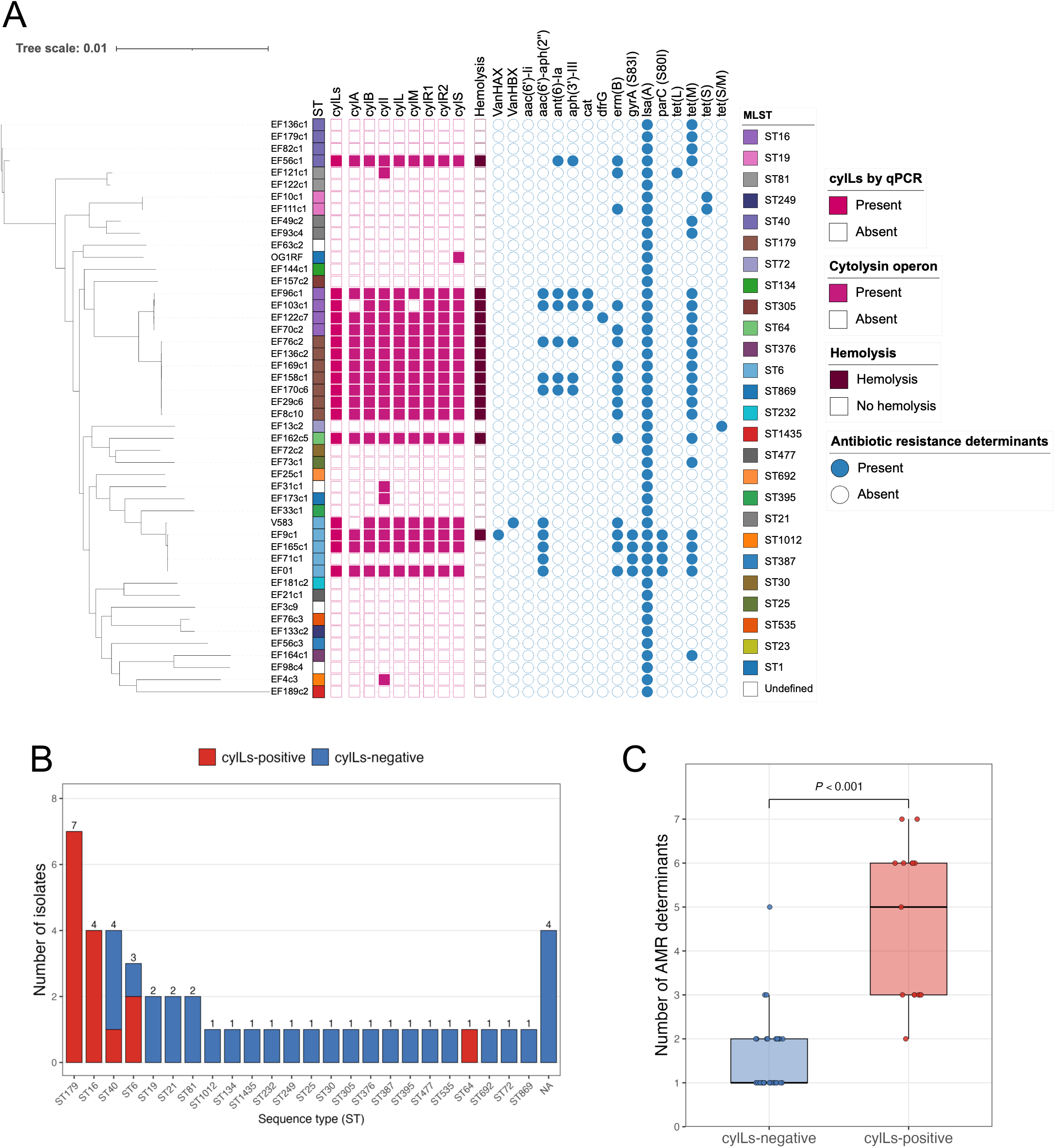
Genomic and phenotypic characteristics of clinical *Enterococcus faecalis* isolates. (A) Core-genome maximum-likelihood phylogeny of 45 clinical *E. faecalis* isolates together with the cytolytic clinical isolate EF01 and the laboratory strains OG1RF and V583. The phylogeny was generated from a core-genome alignment and annotated with multilocus sequence type (MLST), *cylLs* detection by colony qPCR, presence or absence of genes within the cytolysin operon identified from whole-genome sequencing data by ABRicate, beta-hemolytic activity, and antimicrobial resistance (AMR) determinants identified by StarAMR. Filled symbols indicate detection/presence of the corresponding feature and open symbols indicate absence. (B) Distribution of sequence types among the 45 clinical isolates. Red and blue portions of the bars indicate *cylLs*-positive (n = 15) and *cylLs*-negative (n = 30) isolates, respectively. NA indicates isolates for which a sequence type could not be assigned. (C) Comparison of AMR determinant counts between *cylLs*-positive and *cylLs*-negative clinical isolates. Each point represents one isolate. Groups were compared using a two-sided Wilcoxon rank-sum test.

### 3.2. ΦEf2.1 and ΦEf2.2 show increased lytic activity against cylLs-positive clinical isolates

We next evaluated the efficiency of plating (EoP) and the adsorption rate of three *E. faecalis* phages, Φ Ef2.1, Φ Ef2.2, and Φ Ef2.3Ev, against the clinical isolate collection and laboratory strains (Fig. 2A, S2A, and S2B). Both ΦEf2.1 and ΦEf2.3Ev are classified within the genus *Kochikohdavirus* and exhibited similar host-range and adsorption profiles except against ST179 isolates (EF76c2, EF136c2, EF169c1, EF158c1, EF170c6, EF29c6, and EF8c10). ΦEf2.1 showed a low adsorption rate and lytic activity against the ST179 isolates. Despite somewhat reduced adsorption of ΦEf2.3Ev to ST179 isolates, ΦEf2.3Ev maintained an EoP ≥0.1 against all ST179 isolates except EF29c6. In contrast, ΦEf2.2 showed the highest EoP values against most ST179 isolates, except for EF8c10. These results suggest that even closely related *Kochikohdavirus* phages can exhibit different lytic activities and adsorption rates against specific *E. faecalis* lineages. Additionally, 35 out of 45 (77.8%) clinical isolates showed an EoP value of at least 0.01 against one or more of the three phages. The remaining 10 isolates had EoP values of less than 0.01 against all tested phages, indicating resistance or poor susceptibility under the tested conditions. When the 45 clinical isolates were stratified by *cylLs* status, ΦEf2.1 and ΦEf2.2 exhibited significantly higher EoP values against *cylLs*-positive isolates than against *cylLs*-negative isolates (Fig. 2B). In contrast, ΦEf2.3Ev did not exhibit a significant difference in EoP between *cylLs*-positive and - negative isolates. To summarize phage susceptibility across the collection, the isolates were classified as sensitive (EoP ≥ 0.1), intermediate (0.01 ≤ EoP < 0.1), or resistant (EoP < 0.01) based on EoP thresholds. The overall proportions of sensitive isolates across all clinical isolates were 37.8% for ΦEf2.1, 40% for ΦEf2.2, and 53.3% for ΦEf2.3Ev (Fig. S2C). Furthermore, when divided into two groups based on the presence or absence of the *cylLs*, 73.3% of *cylLs*-positive isolates showed at least intermediate susceptibility to ΦEf2.1, compared with 36.7% of *cylLs*-negative isolates. Similarly, 66.7% of the *cylLs*-positive strains were sensitive to ΦEf2.2, while only 26.7% of the *cylLs*-negative strains were sensitive. ΦEf2.3Ev exhibited the broadest host range, with EoP ≥0.01 in 50.0% of *cylLs*-negative isolates and 73.3% of *cylLs*-positive isolates. Adsorption efficiency varied substantially by phage and host strain (Fig. 2A, Fig. S2B, and Fig. S2D). Comparing adsorption between *cylLs*-positive and *cylLs*-negative isolates revealed that ΦEf2.2 had significantly higher adsorption to *cylLs*-positive isolates. Although ΦEf2.3Ev adsorbed significantly more to *cylLs*-negative strains than to *cylLs*-positive strains, the median adsorption rate for the positive strains was still 77%. This finding is consistent with the observation that ΦEf2.3Ev exhibited a slightly lower adsorption rate for ST179 strains, which are *cylLs*-positive. (Fig. S2D). The adsorption rate of ΦEf2.1 did not differ significantly by the *cylLs* status. Finally, we investigated the relationship between the adsorption efficiency of each phage and EoP for clinical isolates, EF01, and laboratory strains (Fig. 2D). A substantial fraction of strains supported both high adsorption and productive infection, defined as adsorption ≥80% and EoP ≥0.1. This productive, high-adsorption phenotype was observed in 39.6% of strains for ΦEf2.1, 31.2% for ΦEf2.2, and 43.8% for ΦEf2.3Ev. However, several strains showed low plating efficiency despite efficient phage adsorption. Specifically, 31.2% of strains for ΦEf2.1, 12.5% for ΦEf2.2, and 37.5% for ΦEf2.3Ev showed high adsorption but strong blocking of productive infection (defined as adsorption ≥80% and EoP <0.01). These findings indicate that efficient adsorption alone is insufficient to ensure efficient plaque formation and are consistent with additional constraints acting after adsorption.

**Figure 2.**
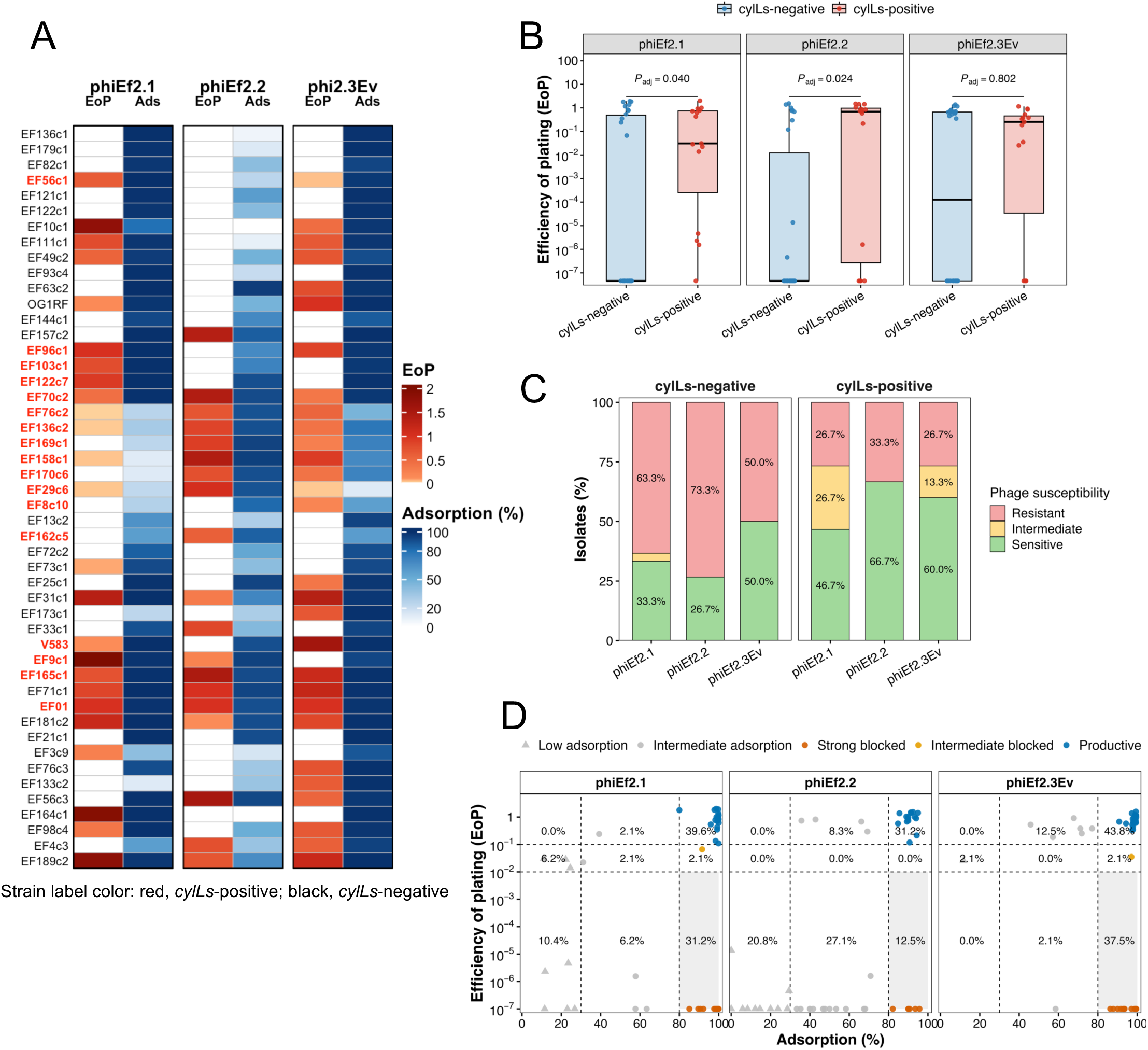
Lytic activity and adsorption profiles of ΦEf2.1, ΦEf2.2, and ΦEf2.3Ev against *Enterococcus faecalis* isolates. (A) Heatmaps showing efficiency of plating (EoP) and adsorption efficiency (Ads) of ΦEf2.1, ΦEf2.2, and ΦEf2.3Ev against the *E. faecalis* isolate collection. Values represent the mean of three replicate measurements. Isolates are ordered according to the core-genome phylogeny shown in Fig. 1A. Red strain names indicate *cylLs*-positive strains. (B) Comparison of EoP values between *cylLs*-negative (n = 30) and *cylLs*-positive (n = 15) clinical isolates. Each point represents the mean EoP from three replicate assays for one isolate. Groups were compared separately for each phage using two-sided Wilcoxon rank-sum tests, and *P* values were adjusted across the three phages using the Benjamini–Hochberg (BH) method. (C) Distribution of phage susceptibility categories among *cylLs*-negative and *cylLs*-positive clinical isolates. Isolates were classified as sensitive (EoP ≥0.1), intermediate (0.01 ≤ EoP <0.1), or resistant (EoP <0.01). Values within bars indicate the percentage of isolates in each category. (D) Relationship between adsorption efficiency and EoP for ΦEf2.1, ΦEf2.2, and ΦEf2.3Ev across all 48 strains analyzed in this study, including the 45 clinical isolates, EF01, OG1RF, and V583. Each point represents the mean of three replicate measurements for one strain. Adsorption was classified as low (<30%), intermediate (30% to <80%), or high (≥80%). High-adsorption infections were further classified as productive (EoP ≥0.1), intermediate blocked (0.01 ≤ EoP <0.1), or strongly blocked (EoP <0.01). Dashed lines indicate the adsorption and EoP thresholds used for classification.

### 3.3. epa variable-region super-patterns are associated with phage adsorption specificity

Given that phage adsorption is one of the predictors of phage host range, we next focused on the *epa* locus, a receptor-associated surface polysaccharide locus implicated in most *E. faecalis* phage infection. The *epa* gene cluster is composed of a highly conserved region extending from *epaA* to *epaR* and a highly variable region subsequent to that point. However, the precise number of patterns that exist remains uncertain. Therefore, to elucidate the diversity of the *epa* variable region, we analyzed 367 complete *E. faecalis* RefSeq genomes obtained from NCBI. The *epa* variable regions were extracted using the highly conserved *epaR* and the EF2163 homolog, which is located immediately downstream of EF2164, the last CDS in the *epa* variable region at V583, as markers. Based on gene composition and sequence similarity, the *epa* variable regions were classified into 15 super-patterns (SP) and named SP001 through SP015 (Figure 3A, Table S2). The architecture-distance tree visualized the relationships among the *epa* variable-region loci, with genomes assigned to the same SP generally occupying neighboring positions (Fig. S3A). The most prevalent groups were SP007, SP005, and SP001, which included 74, 72, and 56 RefSeq genomes, respectively (Fig. S3B). In contrast, SP011 and SP012 contained only two genomes each, and SP013, SP014, and SP015 were singletons. We then extracted *epa* variable region of the clinical isolates and assigned SPs. Assignments were classified as complete when the full-length *epa* variable region was recovered within a single contig, partial when the SP could be inferred from fragmented or partial alignments, and undetermined when insufficient sequence information was available (Fig. S3C). Among the 45 clinical isolates, 24 had complete assignments, 19 had partial assignments, and two could not be assigned reliably to an SP (Table S3). The most frequent SPs were SP005, SP001, SP009, and SP007, whereas SP010 and SP011 were each represented by a single isolate, and no isolates were assigned to SP012 to SP015 (Fig. 3B). Focusing on the *epa* SP distribution of *cylLs*-positive strains, SP009 accounted for the largest number with 8 strains, followed by SP007 with 4 strains (Fig. 3C). Next, we investigated whether *epa* SP is associated with phage adsorption specificity. While ΦEf2.1 and ΦEf2.3Ev exhibited broadly similar adsorption profiles, ΦEf2.3Ev showed a higher adsorption rate than ΦEf2.1 for SP006 and SP009 (Figure 3D). In general, phage ΦEf2.2 demonstrated lower adsorption rates compared to the other two phages. Specifically, its adsorption rate for SP005 was notably lower than that of the other two phages. Conversely, ΦEf2.2 exhibited the highest adsorption rate against SP009. Comparative analysis of the tail-associated genomic regions revealed that most structural genes were conserved between ΦEf2.1 and ΦEf2.3Ev (Fig. S3D). However, one annotated baseplate protein was assigned to a different variable protein group in each phage, indicating divergence in this component of the adsorption apparatus. These data suggest that CDS variations in the *epa* variable region are associated with phage-specific adsorption patterns. Additionally, EoP values among *epa* SPs were compared. The EoP of each phage against isolates of SP009 generally correlated with the adsorption rate (Figure S3E). In contrast, for other strains, substantial variation in EoP values was observed even among isolates classified under the same SP. These results suggest that while the type of the *epa* variable region is associated with phage adsorption specificity, additional factors acting after adsorption likely influence whether productive infection occurs.

**Figure 3.**
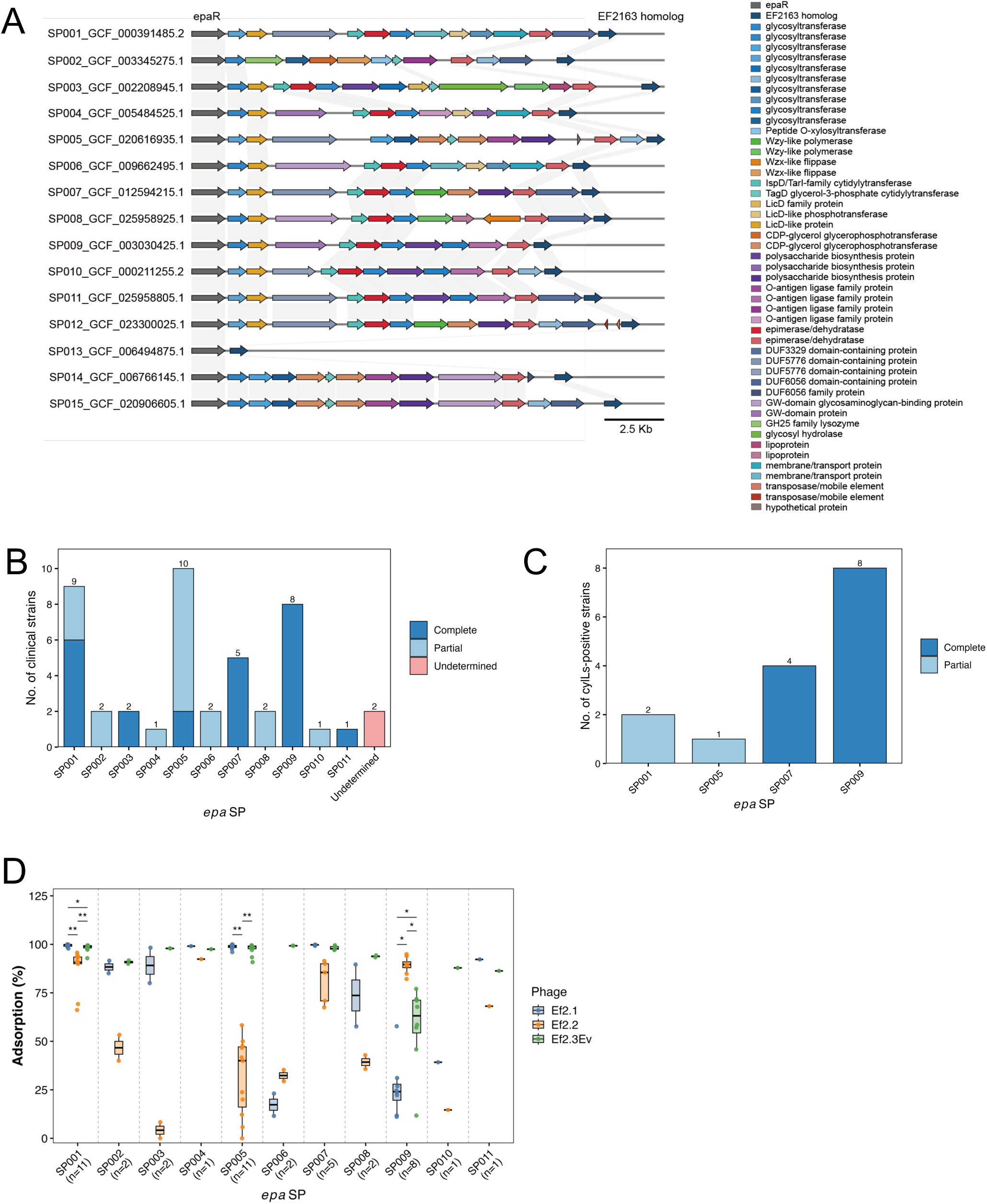
*epa* variable-region super-patterns are associated with phage adsorption specificity. (A) Representative locus architectures of the 15 *epa* variable-region super-patterns (SP001– SP015) identified from 367 quality-controlled complete *E. faecalis* RefSeq genomes. The analyzed locus was defined as the interval from *epaR* (V583 EF2177) to the EF2163 homolog, with the *epa* variable region located downstream of *epaR*. Coding sequences (CDSs) are shown as arrows and colored according to curated functional annotations. Gray links connect CDSs assigned to the same curated orthogroup. (B) Distribution of *epa* variable-region SP assignments among the 45 clinical isolates. Complete assignments indicate recovery and classification of the full *epaR*-to-EF2163 locus; partial assignments indicate SP classification based on recovered partial or terminal *epa* fragments; undetermined isolates could not be reliably assigned to an SP from the available sequence data. (C) Distribution of *epa* variable-region SPs among the 15 *cylLs*-positive clinical isolates. Colors indicate complete and partial assignments, and numbers above bars indicate the number of isolates. (D) Adsorption efficiencies of ΦEf2.1, ΦEf2.2, and ΦEf2.3Ev stratified by *epa* variable-region SP across all 46 strains assigned to an *epa* SP; two strains with undetermined assignments were excluded. Each point represents the mean adsorption efficiency from three replicate assays for one strain. Statistical comparisons were restricted to SPs containing at least three strains. Within each eligible SP, overall differences among the three phages were evaluated using the Friedman test, with measurements from the same bacterial strain treated as matched observations. Pairwise comparisons were performed using paired Wilcoxon signed-rank tests, and the three pairwise *P* values within each SP were adjusted using the BH method. Asterisks indicate BH-adjusted pairwise *P* values: *, P <0.05; **, P <0.01.

### 3.4. CylLs-positive isolates carry more predicted anti-phage defense systems, whereas associations with specific post-adsorption barriers are limited

Since several isolates exhibited efficient phage adsorption but did not support productive infection, we investigated whether predicted anti-phage defense systems were associated with cytolysin status or phage infection outcome. Anti-phage defense systems were identified from the isolate genomes using both DefenseFinder and PADLOC (Fig. S4A, S4B, and Table S4). We first compared the total number of predicted defense system types between the 17 *cylLs*-positive and 31 *cylLs*-negative strains, including all clinical isolates, EF01, V583, and OG1RF. Both prediction tools identified a higher defense-system burden in the *cylLs*-positive group (Fig. 4A). We next examined whether individual defense systems were associated with the outcome of phage infection after efficient adsorption. Given the recent report that the pTEF2-encoded type IV restriction enzyme TIV-RE (EF_B0059) restricts phage replication in *E. faecalis*, a targeted homology search was performed for EF_B0059 in the clinical isolate genomes (Bullen et al., 2024). In this investigation, no homolog of EF_B0059 that satisfied both the predefined sequence identity and coverage thresholds was identified among the 45 clinical isolates. Next, for each phage, isolates with adsorption rates ≥80% were classified as supporting productive infection when EoP was ≥0.1 or as showing strong post-adsorption blocking when EoP was <0.01 (Fig. 2D), and defense-system prevalence was then compared between these two groups using Fisher’s exact tests (Fig. 4B). The analyses identified several defense systems with nominal enrichment in either the strongly blocked or productive-infection groups. Systems enriched among strongly blocked isolates included multiple restriction-modification and abortive infection-associated systems, although no individual defense system remained significantly associated with infection outcome after correction for multiple testing. Nevertheless, AbiD showed the strongest nominal association with strong blocking of ΦEf2.2 infection in both prediction pipelines (Fig. 4B). We therefore compared ΦEf2.2 EoP values between AbiD-positive and AbiD-negative isolates, while also examining the corresponding comparisons for ΦEf2.1 and ΦEf2.3Ev (Fig. 4C). Using DefenseFinder annotations, AbiD-positive isolates tended to show lower ΦEf2.2 EoP values than AbiD-negative isolates, but the difference did not reach statistical significance (*P* = 0.076). No significant differences were observed for ΦEf2.1 (*P* = 0.57) or ΦEf2.3Ev (*P* = 0.23). Using PADLOC annotations, AbiD-positive isolates showed significantly lower ΦEf2.2 EoP values than AbiD-negative isolates (*P* = 0.0066), whereas no significant differences were observed for ΦEf2.1 (*P* = 0.55) or ΦEf2.3Ev (*P* = 0.85).

**Figure 4.**
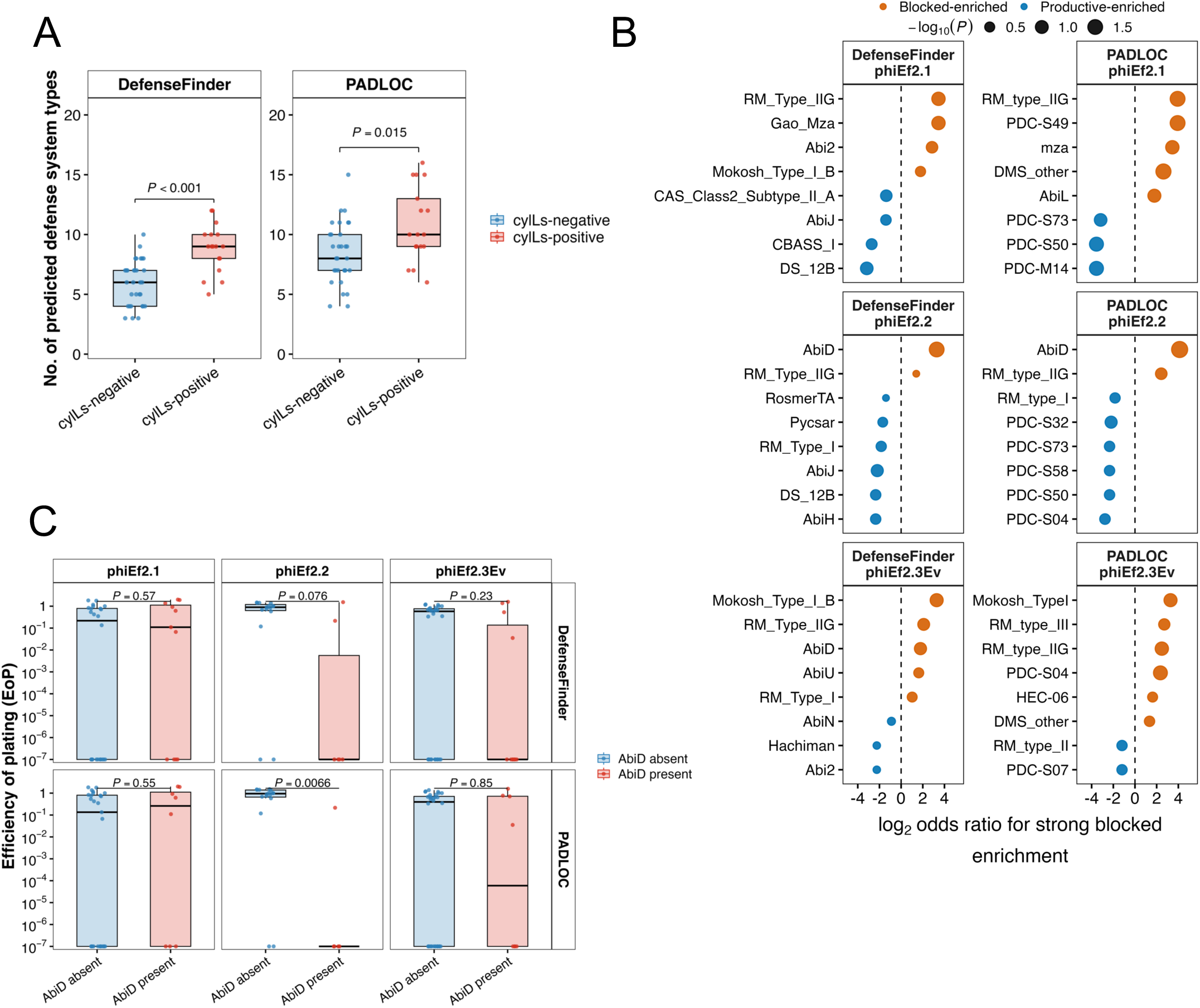
Predicted anti-phage defense systems and post-adsorption restriction of phage infection. (A) Number of predicted anti-phage defense-system types among all 48 *E. faecalis* strains analyzed in this study, stratified by *cylLs* status. Defense systems were independently predicted using DefenseFinder (left) and PADLOC (right). Groups were compared using two-sided Wilcoxon rank-sum tests; *P* values are shown above the comparisons. (B) Association between individual predicted anti-phage defense systems and post-adsorption infection outcome. Analyses were restricted to strain–phage combinations with adsorption ≥80%. For each predicted defense system, its presence or absence was compared between productive and strongly blocked infections using Fisher’s exact test. Productive infection was defined as EoP ≥0.1 and strong post-adsorption blocking as EoP <0.01. For visualization, odds ratios were calculated using a Haldane–Anscombe correction of 0.5 to all cells, and plotted on a log2 scale; positive values indicate enrichment among strongly blocked infections and negative values indicate enrichment among productive infections. Orange and blue points indicate blocked-enriched and productive-enriched systems, respectively, and point size represents −log10 of the nominal *P* value. *P* values were adjusted using the BH method independently within each phage and prediction pipeline. For visualization, only informative defense systems present in at least two of the compared strains were considered, and the eight systems with the smallest nominal P values within each phage–prediction-pipeline combination are shown. No individual defense system remained statistically significant after BH correction. (C) Comparison of EoP values between AbiD-negative and AbiD-positive strains according to DefenseFinder (top row) and PADLOC (bottom row) annotations. Comparisons are shown separately for ΦEf2.1, ΦEf2.2, and ΦEf2.3Ev. Each point represents the mean EoP from three replicate assays for one strain. Groups were compared using two-sided Wilcoxon rank-sum tests.

## 4. Discussion

In this study, we characterized the susceptibility of clinical *E. faecalis* isolates against three phages and investigated bacterial and phage factors associated with their host range. The three phages collectively infected a large part of the collection of clinical isolates: 35 out of 45 isolates (77.8%) showed an EoP of at least 0.01 against at least one phage. ΦEf2.3Ev exhibited the broadest overall host range, whereas ΦEf2.1 and ΦEf2.2 showed preferential activity against *cylLs*-positive isolates. We further identified associations between *epa* variable-region architecture and phage adsorption, while substantial discordance between adsorption and EoP indicated that post-adsorption factors also contribute to productive infection.

The preferential activity of ΦEf2.1 and ΦEf2.2 against *cylLs*-positive isolates may be particularly relevant because these isolates carried a substantially greater burden of antimicrobial resistance determinants. Although antibiotics have been evaluated in patients with alcohol-associated hepatitis, a consistent survival benefit has not been demonstrated. In a randomized trial, prophylactic amoxicillin-clavulanate reduced bacterial infections but did not improve 60-day survival in patients with severe alcohol-associated hepatitis receiving prednisolone (Louvet et al., 2023). Randomized studies of the gut-directed antibiotic rifaximin have similarly failed to demonstrate a significant survival benefit (Kimer et al., 2022; Song et al., 2025). In addition, preclinical studies further suggest that the specificity of microbiota-targeted treatment may be important. In ethanol-fed, gastric acid-suppressed mice, non-absorbable vancomycin and bacitracin reduced intestinal *E. faecalis* but paradoxically exacerbated alcohol-associated liver disease, accompanied by expansion and translocation of other potentially pathogenic bacteria (Raya Tonetti et al., 2026). In contrast, phages selectively targeting cytolysin-positive *E. faecalis* prevented ethanol-induced liver disease in humanized mice colonized with fecal microbiota from patients with alcohol-associated hepatitis (Duan et al., 2019). Together, these findings suggest that selective elimination of a disease-associated pathobiont may offer advantages over broad perturbation of the intestinal microbiota. Although our data do not establish that phage therapy is superior to antibiotics, the increased AMR burden of *cylLs*-positive isolates, together with the preferential activity of ΦEf2.1 and ΦEf2.2 against these strains, provides a rationale for further development of phage-based strategies targeting cytolysin-positive *E. faecalis*. However, because *cylLs*-positive isolates were concentrated within a limited number of sequence types, the observed association between *cylLs* status and phage susceptibility may partly reflect lineage-associated genomic features, including variation in *epa* architecture, rather than a causal effect of *cylLs* itself. An additional potential advantage of phage therapy is that acquisition of resistance to these phages may impose a fitness cost on *E. faecalis*. Our previous work using the cytolytic clinical isolate EF01 demonstrated that variants selected during exposure to ΦEf2.2 acquired alterations affecting Epa biosynthesis, including mutation of *epaR* and loss of *galE*, and showed reduced intestinal colonization, decreased adherence to epithelial cells, and increased susceptibility to bile salts (Fujiki et al., 2025). Thus, although the emergence of phage resistance is an important concern for phage therapy, at least some resistance mechanisms against these phages may reduce bacterial fitness or virulence-associated phenotypes. This possibility is particularly relevant for microbiota-targeted therapies, in which selection for a less fit phage-resistant population may still produce a favorable biological outcome.

A major finding of this study was the association between *epa* variable-region architecture and phage adsorption. Epa consists of a relatively conserved rhamnopolysaccharide backbone decorated with strain-variable, surface-exposed structures encoded by genes downstream of *epaR* (Guerardel et al., 2020). Previous comparative genomic studies have established that this downstream region is highly diverse among *E. faecalis* strains (Palmer et al., 2012). Furthermore, previous genetic studies have shown that disruption of Epa biosynthesis can reduce phage adsorption or confer resistance (Chatterjee et al., 2020, 2019; Ho et al., 2018), and deletion of the V583 *epa* variable region abolished infection by SHEF13, with susceptibility restored by complementation (Alrafaie et al., 2024). However, this diversity has not, to our knowledge, been systematically organized into recurrent architecture-defined groups and directly linked to quantitative phage adsorption phenotypes. Extending these observations to naturally occurring diversity, we classified 367 complete *E. faecalis* RefSeq *epa* variable regions into 15 architecture-based SPs and found distinct phage-specific adsorption patterns. ΦEf2.2 showed particularly strong SP dependence, with low adsorption to SP002, SP003, and SP005 but high adsorption to SP001 and SP009, suggesting that the composition or accessibility of Epa surface decorations can markedly influence adsorption. A recent study also associated *Kochikohdavirus* adsorption with *licD*, which is implicated in Epa decoration (Awad et al., 2026). However, in our clinical isolates, the V583 EF2175 *licD* homolog was present in all represented SPs except SP002, indicating that the presence of EF2175 alone does not explain the observed adsorption patterns (Figure 3A). Rather, LicD-dependent modification may represent one component of the broader Epa decoration architecture that collectively determines the bacterial surface available for phage recognition. A limitation of the *epa* architecture analysis is that SP assignments were based on short-read genome assemblies. Although most clinical isolates could be assigned to an SP, nearly half were classified from partial or terminal *epa* fragments, which may have reduced assignment resolution.

The contrasting adsorption profiles of the two closely related *Kochikohdavirus* phages are consistent with contributions from both bacterial surface variation and differences in phage receptor-recognition machinery. ΦEf2.1 showed markedly reduced adsorption to SP006 and SP009, whereas ΦEf2.3Ev maintained relatively high adsorption across most SPs. Most tail-associated genes were conserved between the two phages, whereas an annotated baseplate protein differed (Fig. S3D), making this protein a candidate determinant of their distinct adsorption specificities. This possibility is supported by studies of other Gram-positive phages in which baseplate-associated receptor-binding proteins directly determine recognition of cell-wall glycopolymers; for example, the baseplate RBP Gp45 of *Staphylococcus aureus* phage Φ11 mediates adsorption to glycosylated wall teichoic acid (Li et al., 2016), while alteration of a baseplate-associated RBP in the Twort-like phage ΦSA012 changes recognition of specific WTA decorations (Takeuchi et al., 2016). Thus, even closely related *Kochikohdavirus* phages may differ in their recognition of, or tolerance to, particular Epa surface structures. The broad adsorption profile of ΦEf2.3Ev raises the possibility that it recognizes an Epa-associated feature conserved across diverse SPs. Although the conserved rhamnan backbone is one potential candidate, structural analysis of V583 indicates that this backbone is largely embedded within the cell wall beneath the surface-exposed decorations (Guerardel et al., 2020). Therefore, ΦEf2.3Ev may instead recognize a conserved feature shared among different Epa decorations or another surface receptor whose accessibility is modulated by Epa.

Interestingly, *cylLs*-positive isolates carried significantly more predicted anti-phage defense system types than *cylLs*-negative isolates, despite being more susceptible to ΦEf2.1 and ΦEf2.2. This contrast suggests that the number of predicted defense systems does not directly reflect the level of protection against a particular phage. Individual defense systems can exhibit strong phage specificity, and combinations of systems can act synergistically (Wu et al., 2024). Moreover, the presence of a predicted defense locus does not necessarily indicate that it is intact, expressed, or effective against the phages tested. Phage-encoded counter-defense mechanisms may further uncouple the presence of a bacterial defense system from the observed infection phenotype. A recent example in *E. faecalis* is the type IV restriction enzyme TIV-RE, which restricts phage phi47 but can be neutralized by the phage-encoded inhibitor TifA (Bullen et al., 2024). We did not detect this TIV-RE in our clinical isolate collection, indicating that this specific mechanism does not explain the post-adsorption restriction observed here. AbiD showed a nominal association with reduced ΦEf2.2 productivity, although this association was prediction-tool dependent and did not remain significant after correction for multiple testing. AbiD belongs to a family of abortive infection systems originally characterized in *Lactococcus lactis*, where it reduces phage propagation without substantially affecting adsorption, consistent with a post-adsorption mode of defense (McLandsborough et al., 1995). To our knowledge, AbiD-mediated phage defense has not been experimentally demonstrated in *E. faecalis*. We therefore consider AbiD a candidate factor for future functional validation rather than a demonstrated mechanism of ΦEf2.2 resistance.

The co-enrichment of AMR determinants and predicted anti-phage defense systems in *cylLs*-positive isolates may reflect lineage-associated differences in accessory-genome composition, including histories of acquisition and retention of mobile genetic elements (MGEs), rather than a direct relationship with *cylLs*. The cytolysin locus itself can be carried on the pheromone-responsive plasmid pAD1 or within a chromosomal pathogenicity island (Gilmore et al., 1994; Shankar et al., 2002). Anti-phage defense systems can themselves be encoded by MGEs, and their accumulation may therefore partly reflect the acquisition and retention of accessory genetic elements rather than simply stronger host immunity (Doron et al., 2018; Makarova et al., 2011; Rocha and Bikard, 2022; Vassallo et al., 2022). Importantly, accumulation of multiple defense systems does not necessarily translate into broad resistance across phages, because their effects depend on ecological context and on the specific defense–MGE–phage combinations involved (Clabby et al., 2025). Against this background, one possible interpretation of our findings is that some *cylLs*-positive lineages possess genomic backgrounds compatible with the acquisition and retention of accessory genetic elements. We speculate that aspects of such a background may also reduce effective restriction of certain phage genomes after entry, thereby allowing productive infection to proceed. However, the enrichment of AMR determinants and predicted defense systems observed here provides only indirect evidence of such an accessory-genome background and does not directly measure MGE acquisition or retention. Thus, the proposed link between permissiveness to accessory genetic elements and phage susceptibility remains a lineage-associated hypothesis rather than evidence that *cylLs* itself promotes either horizontal gene acquisition or phage susceptibility.

Conversely, the relatively low number of currently predicted defense systems in *cylLs*-negative isolates does not exclude the presence of potent antiviral mechanisms. Functional screening has identified multiple anti-phage systems that were missed by conventional genome-context-based computational searches (Vassallo et al., 2022), indicating that the currently annotated bacterial defense systems remain incomplete. Therefore, some *cylLs-*negative isolates that retained efficient adsorption but showed low EoP to all three phages may harbor uncharacterized defense mechanisms or other post-adsorption barriers. These resistant clinical isolates may provide useful candidates for future functional genetic screens to identify additional *E. faecalis* anti-phage mechanisms.

## 5. Conclusion

ΦEf2.1, ΦEf2.2, and ΦEf2.3Ev collectively displayed activity against approximately 78% of the clinical *E. faecalis* isolates examined, supporting further evaluation of these phages as candidate components of a phage cocktail for targeting cytolysin-positive *E. faecalis*. The preferential activity of ΦEf2.1 and ΦEf2.2 against *cylLs*-positive isolates with a higher predicted AMR determinant burden, together with previously described fitness costs associated with some phage-resistance mechanisms, provides additional rationale for their therapeutic development. Our results further identify *epa* variable-region architecture as an important correlate of phage adsorption, while highlighting that efficient infection is additionally constrained by post-adsorption factors. Although *cylLs*-positive strains carried more predicted antiphage defense-system types, they remained relatively susceptible to ΦEf2.1 and ΦEf2.2, suggesting that a greater number of predicted defense systems does not necessarily correspond to stronger protection against a given phage. The contributions of individual defense systems to phage susceptibility remain unresolved. Together, these findings link naturally occurring surface-associated genomic variation to phage adsorption and highlight additional constraints on efficient infection, providing a basis for further evaluation of phages targeting clinically relevant *E. faecalis* populations.

## CRediT authorship contribution statement

**Tomohiro Nakamura:** Conceptualization, Methodology, Investigation, Formal analysis, Data curation, Visualization, Writing – original draft, Writing – review & editing. **Jumpei Fujiki:** Conceptualization, Writing – review & editing. **AlcHepNet Investigators**: Investigation, Resources, Data curation, Writing – review & editing. **Bernd Schnabl:** Conceptualization, Resources, Supervision, Project administration, Funding acquisition, Writing – review & editing.

## Ethics approval and consent to participate

For the AlcHepNet consortium (NCT04072822), ethical oversight was provided centrally by the Western Institutional Review Board (WIRB) (Reference No. 20190963), which served as the central IRB for all participating clinical sites. Written informed consent was obtained from all participants.

## Declaration of generative AI and AI-assisted technologies in the manuscript preparation process

During the preparation of this work, the authors used ChatGPT-5.5 and ChatGPT-5.6 Sol in order to assist with manuscript organization and/or improve language and readability. After using these tools, the authors reviewed and edited the content as needed and take full responsibility for the content of the publication.

## Data availability

All whole-genome sequencing data generated in this study are publicly available through the International Nucleotide Sequence Database Collaboration (INSDC). Draft genome assemblies and associated annotations were deposited in DDBJ/ENA/GenBank under BioProject accession number PRJDB42478, and the corresponding raw sequencing reads were deposited in the DDBJ Sequence Read Archive (DRA). Strain-specific WGS accession numbers, BioSample accession numbers, and DRA Run accession numbers are listed in Table S1.

## Supplementary material

Supplementary material associated with this article includes Supplementary Figures S1–S4, Supplementary Tables S1–S4, and Roster for Authorship from AlcHepNet Investigators.

**Figure S1.** Hemolytic activity and antimicrobial resistance profiles of clinical *Enterococcus faecalis* isolates.

**Figure S2.** Distribution of phage EoP, adsorption, and susceptibility phenotypes among clinical *Enterococcus faecalis* isolates.

**Figure S3.** Architecture-based classification of *epa* variable regions and associated phage phenotypes.

**Figure S4.** Phylogenetic distribution of predicted antiphage defense systems in the *Enterococcus faecalis* strain collection.

**Table S1.** Genomic, cytolysin-associated, hemolytic, and antimicrobial resistance characteristics of Enterococcus faecalis strains used in this study.

**Table S2.** *epa* variable-region architecture classification of 367 complete Enterococcus faecalis RefSeq genomes.

**Table S3.** *epa* variable-region SP assignments of clinical and reference Enterococcus faecalis strains.

**Table S4.** Predicted antiphage defense systems identified in Enterococcus faecalis strains using DefenseFinder and PADLOC.

## Roster for Authorship from AlcHepNet Investigators

**Indiana University School of Medicine, Indianapolis, IN:** Naga Chalasani, MD; Ojo Olawale, MD; Lauren Nephew, MD; Raj Vuppalanchi, MD; Niha Samala, MD; Lindsey Yoder, PA; Suthat Liangpunsakul, MD, MPH, Caitlin Snoeberger, RN; Gail Wind, RN.

**Mayo Clinic, Rochester, Minnesota:** Vijay H. Shah, MD; Douglas A. Simonetto, MD; Patrick Kamath, MD; Victor Karpyak, MD; Amy Olofson, RN.

**Cleveland Clinic, Cleveland, Ohio:** Srinivasan Dasarathy, MD; Nicole Welch, MD; Annette Bellar; Amy Attaway, MD; Jaividhya Dasarathy, MD (MetroHealth, Cleveland, Ohio); Ashley Growley; David Streem, MD; Laura E. Nagy, PhD

**University of Texas Southwestern, Dallas, Texas:** Mack C. Mitchell, MD; Thomas Kerr, MD; Thomas Cotter, MD; Jacquelyn O’Leary, MD; Sherwood Brown, MD; Indira Bhavsar-Burke, MD

**Virginia Commonwealth University, Richmond, Virginia:** Arun J. Sanyal, MD, MBBS; Sara O’Connor, RN; Velimir Luketic, MD; Amon Asgharpour, MD; Juan Pablo Arab MD; Richard K. Sterling MD, MSc; F Gerard Moeller, MD

**University of Louisville, Louisville, Kentucky:** Craig J. McClain, MD; Ashwani K. Singal, MD; Christopher Stewart, MD; Vatsalya Vatsalya, MD, PhD; (Matt Cave, MD; Luis Marsano, MD; Ashutosh Barve, MD, PhD; and Jane Frimodig, PhD-Collaborators)

**University of California at San Diego:** Bernd Schnabl, MD

**Data Coordinating Center at Indiana University:** Wanzhu Tu, PhD; Samer Gawrieh, MD; Qing Tang, MS; Carla Kettler, MS; Jing Su, PhD; Yang Li, PhD; Savannah Yarnelle; Yunpeng Yu, MS; Tae-Hwi L. Schwantes-An, PhD

## References

Alrafaie, A.M., Pyrzanowska, K., Smith, E.M., Partridge, D.G., Rafferty, J., Mesnage, S., Shepherd, J., Stafford, G.P., 2024. A diverse set of Enterococcus-infecting phage provides insight into phage host-range determinants. Virus Res. 347, 199426. 10.1016/j.virusres.2024.199426

Al-Zubidi, M., Widziolek, M., Court, E.K., Gains, A.F., Smith, R.E., Ansbro, K., Alrafaie, A., Evans, C., Murdoch, C., Mesnage, S., Douglas, C.W.I., Rawlinson, A., Stafford, G.P., 2019. Identification of Novel Bacteriophages with Therapeutic Potential That Target Enterococcus faecalis. Infect. Immun. 87, e00512–19. 10.1128/IAI.00512-19

Awad, M., Bouras, G., Feizi, S., Grigson, S.R., Wormald, P.-J., Psaltis, A.J., Vreugde, S., 2026. LicD-mediated cell wall decoration governs phage sensitivity in Enterococcus faecalis clinical isolates. Microbiol. Res. 302, 128341. 10.1016/j.micres.2025.128341

Bernheim, A., Sorek, R., 2020. The pan-immune system of bacteria: antiviral defence as a community resource. Nat. Rev. Microbiol. 18, 113–119. 10.1038/s41579-019-0278-2

Bharat, A., Petkau, A., Avery, B.P., Chen, J.C., Folster, J.P., Carson, C.A., Kearney, A., Nadon, C., Mabon, P., Thiessen, J., Alexander, D.C., Allen, V., El Bailey, S., Bekal, S., German, G.J., Haldane, D., Hoang, L., Chui, L., Minion, J., Zahariadis, G., Van Domselaar, G., Reid-Smith, R.J., Mulvey, M.R., 2022. Correlation between Phenotypic and In Silico Detection of Antimicrobial Resistance in Salmonella enterica in Canada Using Staramr. Microorganisms 10, 292. 10.3390/microorganisms10020292

Bolger, A.M., Lohse, M., Usadel, B., 2014. Trimmomatic: a flexible trimmer for Illumina sequence data. Bioinformatics 30, 2114–2120. 10.1093/bioinformatics/btu170

Bouras, G., Grigson, S.R., Mirdita, M., Heinzinger, M., Papudeshi, B., Mallawaarachchi, V., Green, R., Kim, R.S., Mihalia, V., Psaltis, A.J., Wormald, P.-J., Vreugde, S., Steinegger, M., Edwards, R.A., 2026. Protein structure-informed bacteriophage genome annotation with Phold. Nucleic Acids Res. 54, gkaf1448. 10.1093/nar/gkaf1448

Bouras, G., Nepal, R., Houtak, G., Psaltis, A.J., Wormald, P.-J., Vreugde, S., 2023. Pharokka: a fast scalable bacteriophage annotation tool. Bioinformatics 39, btac776. 10.1093/bioinformatics/btac776

Bullen, N.P., Johnson, C.N., Andersen, S.E., Arya, G., Marotta, S.R., Lee, Y.-J., Weigele, P.R., Whitney, J.C., Duerkop, B.A., 2024. An enterococcal phage protein inhibits type IV restriction enzymes involved in antiphage defense. Nat. Commun. 15, 6955. 10.1038/s41467-024-51346-1

Cabré, N., Yang, Y., Wang, Y., Schnabl, B., 2023. Development of a Quantitative PCR Method for Detecting Enterococcus faecalis Cytolysin in Human Stool Samples. Methods Protoc. 6, 107. 10.3390/mps6060107

Camacho, C., Coulouris, G., Avagyan, V., Ma, N., Papadopoulos, J., Bealer, K., Madden, T.L., 2009. BLAST+: architecture and applications. BMC Bioinformatics 10, 421. 10.1186/1471-2105-10-421

Chatterjee, A., Johnson, C.N., Luong, P., Hullahalli, K., McBride, S.W., Schubert, A.M., Palmer, K.L., Carlson, P.E.J., Duerkop, B.A., 2019. Bacteriophage resistance alters antibiotic-mediated intestinal expansion of enterococci. Infect. Immun. 87, e00085–19. 10.1128/IAI.00085-19

Chatterjee, A., Willett, J.L.E., Nguyen, U.T., Monogue, B., Palmer, K.L., Dunny, G.M., Duerkop, B.A., 2020. Parallel Genomics Uncover Novel Enterococcal-Bacteriophage Interactions. mBio 11, e03120–19. 10.1128/mBio.03120-19

Chen, L., Zheng, D., Liu, B., Yang, J., Jin, Q., 2016. VFDB 2016: hierarchical and refined dataset for big data analysis—10 years on. Nucleic Acids Res. 44, D694–D697. 10.1093/nar/gkv1239

Clabby, T., Tesson, F., Gaborieau, B., Bernheim, A., 2025. Why do bacteria accumulate antiphage defence systems? Philosophical Transactions of the Royal Society B: Biological Sciences 380, 20240082. 10.1098/rstb.2024.0082

Cock, P.J.A., Antao, T., Chang, J.T., Chapman, B.A., Cox, C.J., Dalke, A., Friedberg, I., Hamelryck, T., Kauff, F., Wilczynski, B., de Hoon, M.J.L., 2009. Biopython: freely available Python tools for computational molecular biology and bioinformatics. Bioinformatics 25, 1422–1423. 10.1093/bioinformatics/btp163

Dedrick, R.M., Guerrero-Bustamante, C.A., Garlena, R.A., Russell, D.A., Ford, K., Harris, K., Gilmour, K.C., Soothill, J., Jacobs-Sera, D., Schooley, R.T., Hatfull, G.F., Spencer, H., 2019. Engineered bacteriophages for treatment of a patient with a disseminated drug-resistant Mycobacterium abscessus. Nat. Med. 25, 730–733. 10.1038/s41591-019-0437-z

Doron, S., Melamed, S., Ofir, G., Leavitt, A., Lopatina, A., Keren, M., Amitai, G., Sorek, R., 2018. Systematic discovery of antiphage defense systems in the microbial pangenome. Science. 359, eaar4120. 10.1126/science.aar4120

Duan, Y., Llorente, C., Lang, S., Brandl, K., Chu, H., Jiang, L., White, R.C., Clarke, T.H., Nguyen, K., Torralba, M., Shao, Y., Liu, J., Hernandez-Morales, A., Lessor, L., Rahman, I.R., Miyamoto, Y., Ly, M., Gao, B., Sun, W., Kiesel, R., Hutmacher, F., Lee, S., Ventura-Cots, M., Bosques-Padilla, F., Verna, E.C., Abraldes, J.G., Brown, R.S., Vargas, V., Altamirano, J., Caballería, J., Shawcross, D.L., Ho, S.B., Louvet, A., Lucey, M.R., Mathurin, P., Garcia-Tsao, G., Bataller, R., Tu, X.M., Eckmann, L., van der Donk, W.A., Young, R., Lawley, T.D., Stärkel, P., Pride, D., Fouts, D.E., Schnabl, B., 2019. Bacteriophage targeting of gut bacterium attenuates alcoholic liver disease. Nature 575, 505–511. 10.1038/s41586-019-1742-x

Egorov, A.A., Atkinson, G.C., 2025. LoVis4u: a locus visualization tool for comparative genomics and coverage profiles. NAR Genom. Bioinform. 7, lqaf009. 10.1093/nargab/lqaf009

Federici, S., Kredo-Russo, S., Valdés-Mas, R., Kviatcovsky, D., Weinstock, E., Matiuhin, Y., Silberberg, Y., Atarashi, K., Furuichi, M., Oka, A., Liu, B., Fibelman, M., Weiner, I.N., Khabra, E., Cullin, N., Ben-Yishai, N., Inbar, D., Ben-David, H., Nicenboim, J., Kowalsman, N., Lieb, W., Kario, E., Cohen, T., Geffen, Y.F., Zelcbuch, L., Cohen, A., Rappo, U., Gahali-Sass, I., Golembo, M., Lev, V., Dori-Bachash, M., Shapiro, H., Moresi, C., Cuevas-Sierra, A., Mohapatra, G., Kern, L., Zheng, D., Nobs, S.P., Suez, J., Stettner, N., Harmelin, A., Zak, N., Puttagunta, S., Bassan, M., Honda, K., Sokol, H., Bang, C., Franke, A., Schramm, C., Maharshak, N., Sartor, R.B., Sorek, R., Elinav, E., 2022. Targeted suppression of human IBD-associated gut microbiota commensals by phage consortia for treatment of intestinal inflammation. Cell 185, 2879–2898.e24. 10.1016/j.cell.2022.07.003

Fujiki, J., Nakamura, T., Kreimeyer, H., Llorente, C., Fouts, D.E., Schnabl, B., 2025. Insertion sequence-mediated phage resistance contributes to attenuated colonization of cytolytic Enterococcus faecalis variants in the gut. Microbiol. Spectr. 13, e0330324. 10.1128/spectrum.03303-24

Fujiki, J., Nakamura, T., Le, T., Kobayashi, H., Morales, S., Pride, D., Schnabl, B., 2026. Evolution-guided engineering of a candidate therapeutic phage accelerates intestinal clearance of Enterococcus faecalis. Preprint in preparation.

Fujiki, J., Nakamura, T., Nakamura, K., Nishida, K., Amano, Y., Watanabe, Y., Gondaira, S., Usui, M., Shimizu, M., Miyanaga, K., Watanabe, S., Iwasaki, T., Kiga, K., Hanawa, T., Higuchi, H., Sawa, T., Tanji, Y., Tamura, Y., Cui, L., Iwano, H., 2022. Biological properties of Staphylococcus virus ΦSA012 for phage therapy. Sci. Rep. 12, 21297. 10.1038/s41598-022-25352-6

Fujiki, J., Yoshida, S., Nakamura, T., Nakamura, K., Amano, Y., Nishida, K., Nishi, K., Sasaki, M., Iwasaki, T., Sawa, H., Komatsuzawa, H., Hijioka, H., Iwano, H., 2021. Novel Virulent Bacteriophage ΦSG005, Which Infects Streptococcus gordonii, Forms a Distinct Clade among Streptococcus Viruses. Viruses 13, 1964. 10.3390/v13101964

Gan, L., Feng, Y., Du, B., Fu, H., Tian, Z., Xue, G., Yan, C., Cui, X., Zhang, R., Cui, J., Zhao, H., Feng, J., Xu, Z., Fan, Z., Fu, T., Du, S., Liu, S., Zhang, Q., Yu, Z., Sun, Y., Yuan, J., 2023. Bacteriophage targeting microbiota alleviates non-alcoholic fatty liver disease induced by high alcohol-producing Klebsiella pneumoniae. Nat. Commun. 14, 3215. 10.1038/s41467-023-39028-w

Gilmore, M.S., Segarra, R.A., Booth, M.C., Bogie, C.P., Hall, L.R., Clewell, D.B., 1994. Genetic structure of the Enterococcus faecalis plasmid pAD1-encoded cytolytic toxin system and its relationship to lantibiotic determinants. J. Bacteriol. 176, 7335–7344. 10.1128/jb.176.23.7335-7344.1994

Guerardel, Y., Sadovskaya, I., Maes, E., Furlan, S., Chapot-Chartier, M.-P., Mesnage, S., Rigottier-Gois, L., Serror, P., 2020. Complete Structure of the Enterococcal Polysaccharide Antigen (EPA) of Vancomycin-Resistant Enterococcus faecalis V583 Reveals that EPA Decorations Are Teichoic Acids Covalently Linked to a Rhamnopolysaccharide Backbone. mBio 11, e00277–20. 10.1128/mBio.00277-20

Guindon, S., Dufayard, J.-F., Lefort, V., Anisimova, M., Hordijk, W., Gascuel, O., 2010. New Algorithms and Methods to Estimate Maximum-Likelihood Phylogenies: Assessing the Performance of PhyML 3.0. Syst. Biol. 59, 307–321. 10.1093/sysbio/syq010

Ho, K., Huo, W., Pas, S., Dao, R., Palmer, K.L., 2018. Loss-of-Function Mutations in epaR Confer Resistance to ϕNPV1 Infection in Enterococcus faecalis OG1RF. Antimicrob. Agents Chemother. 62, 1–11. 10.1128/AAC.00758-18

Hoang, D.T., Chernomor, O., von Haeseler, A., Minh, B.Q., Vinh, L.S., 2018. UFBoot2: Improving the Ultrafast Bootstrap Approximation. Mol. Biol. Evol. 35, 518–522. 10.1093/molbev/msx281

Hsu, B.B., Gibson, T.E., Yeliseyev, V., Liu, Q., Lyon, L., Bry, L., Silver, P.A., Gerber, G.K., 2019. Dynamic Modulation of the Gut Microbiota and Metabolome by Bacteriophages in a Mouse Model. Cell Host Microbe 25, 803–814.e5. 10.1016/j.chom.2019.05.001

Jolley, K.A., Bray, J.E., Maiden, M.C.J., 2018. Open-access bacterial population genomics: BIGSdb software, the PubMLST.org website and their applications. Wellcome Open Res. 3, 124. 10.12688/wellcomeopenres.14826.1

Jophlin, L.L., Singal, A.K., Bataller, R., Wong, R.J., Sauer, B.G., Terrault, N.A., Shah, V.H., 2024. ACG Clinical Guideline: Alcohol-Associated Liver Disease. American Journal of Gastroenterology 119, 30–54. 10.14309/ajg.0000000000002572

Kalyaanamoorthy, S., Minh, B.Q., Wong, T.K.F., von Haeseler, A., Jermiin, L.S., 2017. ModelFinder: fast model selection for accurate phylogenetic estimates. Nat. Methods 14, 587–589. 10.1038/nmeth.4285

Katoh, K., Standley, D.M., 2013. MAFFT Multiple Sequence Alignment Software Version 7: Improvements in Performance and Usability. Mol. Biol. Evol. 30, 772–780. 10.1093/molbev/mst010

Kimer, N., Meldgaard, M., Hamberg, O., Kronborg, T.M., Lund, A.M., Møller, H.J., Bendtsen, F., Ytting, H., 2022. The impact of rifaximin on inflammation and metabolism in alcoholic hepatitis: A randomized clinical trial. PLoS One 17, e0264278. 10.1371/journal.pone.0264278

Kortright, K.E., Chan, B.K., Koff, J.L., Turner, P.E., 2019. Phage Therapy: A Renewed Approach to Combat Antibiotic-Resistant Bacteria. Cell Host Microbe 25, 219–232. 10.1016/j.chom.2019.01.014

Letunic, I., Bork, P., 2024. Interactive Tree of Life (iTOL) v6: recent updates to the phylogenetic tree display and annotation tool. Nucleic Acids Res. 52, W78–W82. 10.1093/nar/gkae268

Li, X., Koç, C., Kühner, P., Stierhof, Y., Krismer, B., Enright, M.C., Penadés, J.R., Wolz, C., Stehle, T., Cambillau, C., Peschel, A., Xia, G., 2016. An essential role for the baseplate protein Gp45 in phage adsorption to Staphylococcus aureus. Sci. Rep. 6, 26455. 10.1038/srep26455

Louvet, A., Labreuche, J., Dao, T., Thévenot, T., Oberti, F., Bureau, C., Paupard, T., Nguyen-Khac, E., Minello, A., Bernard-Chabert, B., Anty, R., Wartel, F., Carbonell, N., Pageaux, G.-P., Hilleret, M.-N., Moirand, R., Nahon, P., Potey, C., Duhamel, A., Mathurin, P., 2023. Effect of Prophylactic Antibiotics on Mortality in Severe Alcohol-Related Hepatitis: A Randomized Clinical Trial. JAMA 329, 1558–1566. 10.1001/jama.2023.4902

Makarova, K.S., Wolf, Y.I., Snir, S., Koonin, E. V, 2011. Defense Islands in Bacterial and Archaeal Genomes and Prediction of Novel Defense Systems. J. Bacteriol. 193, 6039–6056. 10.1128/JB.05535-11

McLandsborough, L.A., Kolaetis, K.M., Requena, T., McKay, L.L., 1995. Cloning and characterization of the abortive infection genetic determinant abiD isolated from pBF61 of Lactococcus lactis subsp. lactis KR5. Appl. Environ. Microbiol. 61, 2023–6. 10.1128/aem.61.5.2023-2026.1995

Palmer, K.L., Godfrey, P., Griggs, A., Kos, V.N., Zucker, J., Desjardins, C., Cerqueira, G., Gevers, D., Walker, S., Wortman, J., Feldgarden, M., Haas, B., Birren, B., Gilmore, M.S., 2012. Comparative Genomics of Enterococci: Variation in Enterococcus faecalis, Clade Structure in E. faecium, and Defining Characteristics of E. gallinarum and E. casseliflavus. mBio 3, e00318–11. 10.1128/mBio.00318-11

Payne, L.J., Meaden, S., Mestre, M.R., Palmer, C., Toro, N., Fineran, P.C., Jackson, S.A., 2022. PADLOC: a web server for the identification of antiviral defence systems in microbial genomes. Nucleic Acids Res. 50, W541–W550. 10.1093/nar/gkac400

Payne, L.J., Todeschini, T.C., Wu, Y., Perry, B.J., Ronson, C.W., Fineran, P.C., Nobrega, F.L., Jackson, S.A., 2021. Identification and classification of antiviral defence systems in bacteria and archaea with PADLOC reveals new system types. Nucleic Acids Res. 49, 10868– 10878. 10.1093/nar/gkab883

Pirnay, J.P., Djebara, S., Steurs, G., Griselain, J., Cochez, C., De Soir, S., Glonti, T., Spiessens, A., Vanden Berghe, E., Green, S., Wagemans, J., Lood, C., Schrevens, E., Chanishvili, N., Kutateladze, M., de Jode, M., Ceyssens, P.J., Draye, J.P., Verbeken, G., De Vos, D., Rose, T., Onsea, J., Van Nieuwenhuyse, B., Soentjens, P., Lavigne, R., Merabishvili, M., Pang, K.W., Metsemakers, W.J., Van der Linden, D., Chatzis, O., Eskenazi, A., Lopez, A., De Voeght, A., Rousseau, A.F., Tilmanne, A., Vens, D., Gérain, J., Layeux, B., Vlieghe, E., Baar, I., Van Ierssel, S., Van Laethem, J., Guiot, J., De Roock, S., Jennes, S., Uyttebroek, S., Van Gerven, L., Hellings, P.W., Dupont, L., Debaveye, Y., Devolder, D., Spriet, I., De Munter, P., Depypere, M., Vanfleteren, M., Cornu, O., Verhulst, S., Boiy, T., Lamote, S., Van Zele, T., Wieërs, G., Courtin, C., Lebeaux, D., Sartre, J., Ferry, T., Laurent, F., Paul, K., Di Luca, M., Gottschlich, S., Tkhilaishvili, T., Cesta, N., Racenis, K., Barbosa, T., López-Cortés, L.E., Tomás, M., Hübner, M., Pham, T.T., Nagtegaal, P., Ten Oever, J., Daniels, J., Loubert, M., Iheb, G., Jones, J., Hall, L., Young, M., Balarjishvili, N., Tediashvili, M., Tong, Y., Rohde, C., Wittmann, J., Hazan, R., Nir-Paz, R., Azeredo, J., Krylov, V., Cameron, D., Pitton, M., Que, Y.A., Resch, G., McCallin, S., Dunne, M., Kilcher, S., 2024. Personalized bacteriophage therapy outcomes for 100 consecutive cases: a multicentre, multinational, retrospective observational study. Nat. Microbiol. 9, 1434– 1453. 10.1038/s41564-024-01705-x

Raya Tonetti, F., Han, H., Fondevila, M.F., Wei, W., Özdirik, B., Bajaj, J.S., Schubert, M.L., Sikaroodi, M., Gillevet, P.M., Lang, S., Demir, M., Rahman, I.R., van der Donk, W.A., Bosques-Padilla, F., Verna, E.C., Abraldes, J.G., Brown, R.S., Vargas, V., Altamirano, J., Caballería, J., Shawcross, D.L., Louvet, A., Lucey, M.R., Mathurin, P., Garcia-Tsao, G., Stärkel, P., Bataller, R., Hsu, C.L., Llorente, C., 2026. Non-absorbable antibiotics worsen alcohol-associated liver disease in gastric acid-suppressed mice. Gut Microbes 18, 2694797. 10.1080/19490976.2026.2694797

Reynolds, D.L., Simpson, E.B., Hille, M.M., 2025. A Method for Demonstrating the Cytolysin/Hemolysin of Enterococcus faecalis Isolates of Poultry Origin. Poultry 4, 11. 10.3390/poultry4010011

Rigottier-Gois, L., Madec, C., Navickas, A., Matos, R.C., Akary-Lepage, E., Mistou, M.-Y., Serror, P., 2015. The Surface Rhamnopolysaccharide Epa of Enterococcus faecalis Is a Key Determinant of Intestinal Colonization. Journal of Infectious Diseases 211, 62–71. 10.1093/infdis/jiu402

Rocha, E.P.C., Bikard, D., 2022. Microbial defenses against mobile genetic elements and viruses: Who defends whom from what? PLoS Biol. 20, e3001514. 10.1371/journal.pbio.3001514

Schooley, R.T., Biswas, B., Gill, J.J., Hernandez-Morales, A., Lancaster, J., Lessor, L., Barr, J.J., Reed, S.L., Rohwer, F., Benler, S., Segall, A.M., Taplitz, R., Smith, D.M., Kerr, K., Kumaraswamy, M., Nizet, V., Lin, L., McCauley, M.D., Strathdee, S.A., Benson, C.A., Pope, R.K., Leroux, B.M., Picel, A.C., Mateczun, A.J., Cilwa, K.E., Regeimbal, J.M., Estrella, L.A., Wolfe, D.M., Henry, M.S., Quinones, J., Salka, S., Bishop-Lilly, K.A., Young, R., Hamilton, T., 2017. Development and Use of Personalized Bacteriophage-Based Therapeutic Cocktails To Treat a Patient with a Disseminated Resistant Acinetobacter baumannii Infection. Antimicrob. Agents Chemother. 61, 1–14. 10.1128/AAC.00954-17

Schwengers, O., Jelonek, L., Dieckmann, M.A., Beyvers, S., Blom, J., Goesmann, A., 2021. Bakta: rapid and standardized annotation of bacterial genomes via alignment-free sequence identification. Microb. Genom. 7, 685. 10.1099/mgen.0.000685

Shankar, N., Baghdayan, A.S., Gilmore, M.S., 2002. Modulation of virulence within a pathogenicity island in vancomycin-resistant Enterococcus faecalis. Nature 417, 746–750. 10.1038/nature00802

Song, D.S., Yang, J.M., Jung, Y.K., Yim, H.J., Kim, H.Y., Kim, C.W., Kim, S.S., Cheong, J.Y., Lee, H.L., Lee, S.W., Yoo, J.-J., Kim, S.G., Kim, Y.S., 2025. Rifaximin treatment in patients with severe alcohol-associated hepatitis: A multicenter, randomized controlled, open-label, pilot trial. Ann. Hepatol. 30, 101749. 10.1016/j.aohep.2024.101749

Steinegger, M., Söding, J., 2017. MMseqs2 enables sensitive protein sequence searching for the analysis of massive data sets. Nat. Biotechnol. 35, 1026–1028. 10.1038/nbt.3988

Takeuchi, I., Osada, K., Azam, A.H., Asakawa, H., Miyanaga, K., Tanji, Y., 2016. The Presence of Two Receptor-Binding Proteins Contributes to the Wide Host Range of Staphylococcal Twort-Like Phages. Appl. Environ. Microbiol. 82, 5763–5774. 10.1128/AEM.01385-16

Tesson, F., Hervé, A., Mordret, E., Touchon, M., D’Humières, C., Cury, J., Bernheim, A., 2022. Systematic and quantitative view of the antiviral arsenal of prokaryotes. Nat. Commun. 13, 2561. 10.1038/s41467-022-30269-9

Tonkin-Hill, G., MacAlasdair, N., Ruis, C., Weimann, A., Horesh, G., Lees, J.A., Gladstone, R.A., Lo, S., Beaudoin, C., Floto, R.A., Frost, S.D.W., Corander, J., Bentley, S.D., Parkhill, J., 2020. Producing polished prokaryotic pangenomes with the Panaroo pipeline. Genome Biol. 21, 180. 10.1186/s13059-020-02090-4

Van Tyne, D., Martin, M., Gilmore, M., 2013. Structure, Function, and Biology of the Enterococcus faecalis Cytolysin. Toxins (Basel). 5, 895–911. 10.3390/toxins5050895

Vassallo, C.N., Doering, C.R., Littlehale, M.L., Teodoro, G.I.C.C., Laub, M.T., 2022. A functional selection reveals previously undetected anti-phage defence systems in the E. coli pangenome. Nat. Microbiol. 7, 1568–1579. 10.1038/s41564-022-01219-4

Wong, T.K.F., Ly-Trong, N., Ren, H., Demotte, P., Baños, H., Roger, A.J., Susko, E., Bielow, C., De Maio, N., Goldman, N., Hahn, M.W., dos Reis, M., Vinh, L.S., Huttley, G., Lanfear, R., Minh, B.Q., 2026. IQ-TREE 3: phylogenomic inference software using complex evolutionary models. Mol. Biol. Evol. 43, msag117. 10.1093/molbev/msag117

Wu, Y., Garushyants, S.K., van den Hurk, A., Aparicio-Maldonado, C., Kushwaha, S.K., King, C.M., Ou, Y., Todeschini, T.C., Clokie, M.R.J., Millard, A.D., Gençay, Y.E., Koonin, E. V., Nobrega, F.L., 2024. Bacterial defense systems exhibit synergistic anti-phage activity. Cell Host Microbe 32, 557–572.e6. 10.1016/j.chom.2024.01.015

