## Supplemental Figures for "Bacteriophage susceptibility in clinical *Enterococcus faecalis* reflects host genomic background and Epa-associated phage adsorption"

### Figure S1

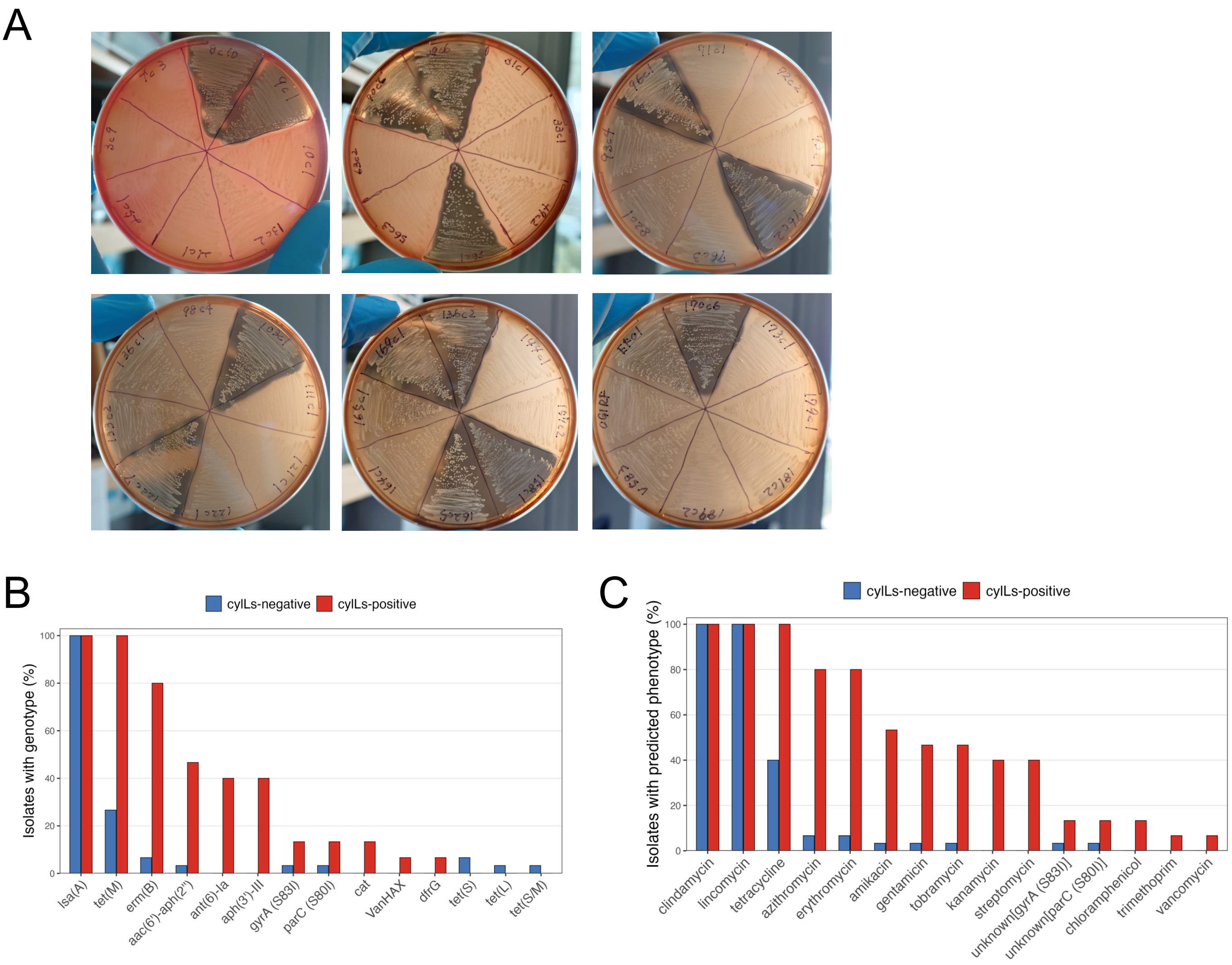


**Figure S1. Hemolytic activity and antimicrobial resistance profiles of clinical *Enterococcus faecalis* isolates.**
(A) Horse blood agar plates used to assess beta-hemolytic activity of *E. faecalis* isolates. Beta-hemolysis was defined as a clear zone surrounding bacterial growth. (B) Prevalence of individual AMR determinants among *cylLs*-negative (blue; n = 30) and *cylLs*-positive (red; n = 15) clinical isolates. Bars indicate the percentage of isolates in each group harboring the indicated determinant. (C) Distribution of antimicrobial resistance phenotypes predicted from AMR determinants identified by StarAMR. Bars indicate the percentage of isolates in each *cylLs* group with the indicated predicted phenotype.

### Figure S2

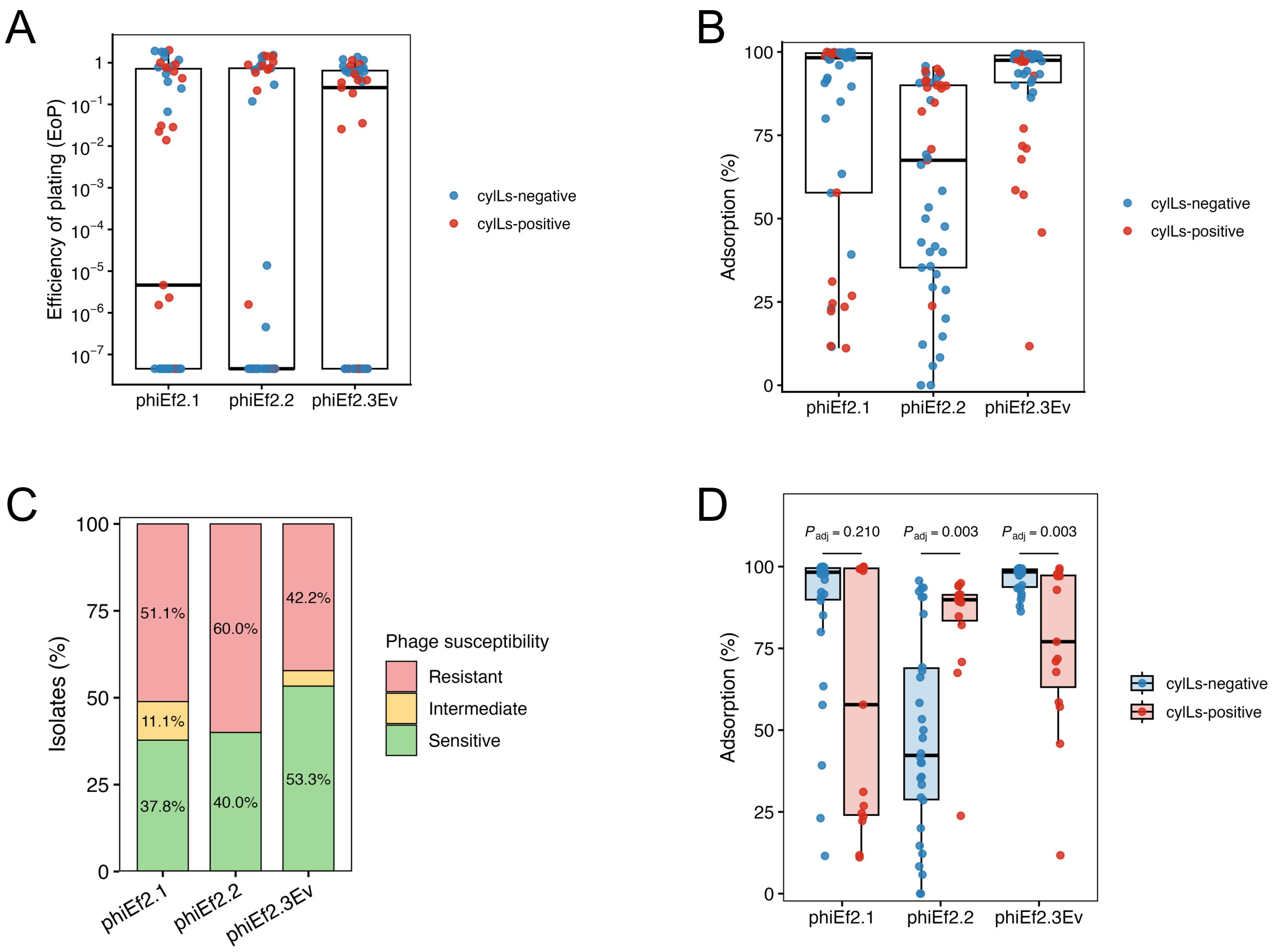


**Figure S2. Distribution of phage EoP, adsorption, and susceptibility phenotypes among clinical *Enterococcus faecalis* isolates.**
(A) Distribution of EoP values for ΦEf2.1, ΦEf2.2, and ΦEf2.3Ev across the 45 clinical isolates. Blue and red points indicate *cylLs*-negative and *cylLs*-positive isolates, respectively. Each point represents the mean of three replicate assays. (B) Distribution of adsorption efficiencies for the three phages across the 45 clinical isolates. Each point represents the mean of three replicate assays. (C) Overall proportions of the 45 clinical isolates classified as sensitive, intermediate, or resistant to each phage. Susceptibility categories were defined as sensitive (EoP ≥ 0.1), intermediate (0.01 ≤ EoP < 0.1), and resistant (EoP < 0.01). Percentages are indicated within the bars. (D) Comparison of adsorption efficiencies between *cylLs*-negative (n = 30) and *cylLs*-positive (n = 15) clinical isolates. Groups were compared separately for each phage using two-sided Wilcoxon rank-sum tests, and *P* values were adjusted across the three phages using the BH method. BH-adjusted *P* values are shown.

### Figure S3

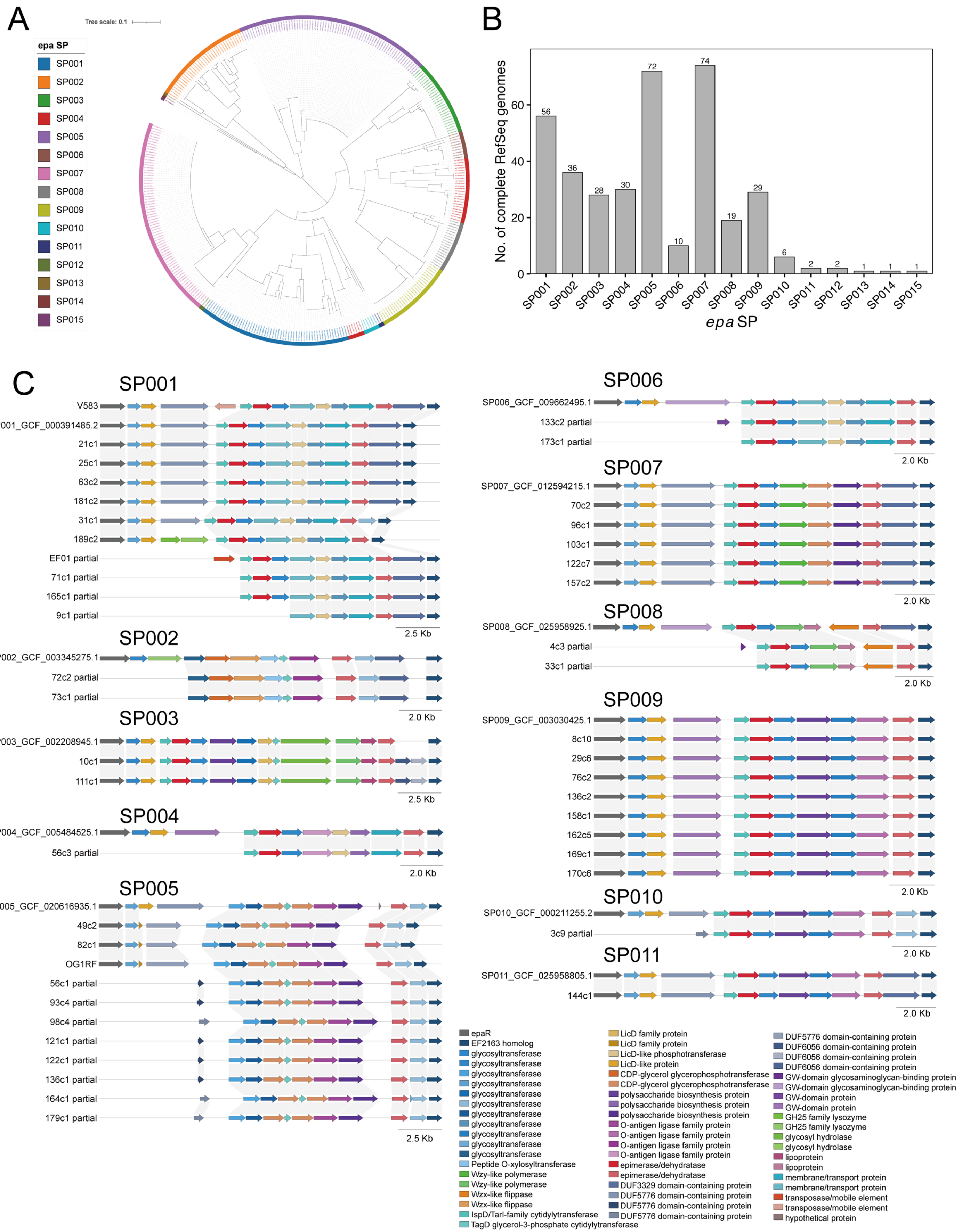

A
Tree scale: 0.1
epa SP
SP001
SP002
SP003
SP004
SP005
SP006
SP007
SP008
SP009
SP010
SP011
SP012
SP013
SP014
SP015
B
No. of complete RefSeq genomes
epa SP
56
36
28
30
72
10
74
19
29
6
2
2
1
1
1
SP001
SP002
SP003
SP004
SP005
SP006
SP007
SP008
SP009
SP010
SP011
SP012
SP013
SP014
SP015
C
SP001
V583
SP001_GCF_000391485.2
21c1
25c1
63c2
181c2
31c1
189c2
EF01 partial
71c1 partial
165c1 partial
9c1 partial
2.5 Kb
SP002
SP002_GCF_003345275.1
72c2 partial
73c1 partial
2.0 Kb
SP003
SP003_GCF_002208945.1
10c1
111c1
2.5 Kb
SP004
SP004_GCF_005484525.1
56c3 partial
2.0 Kb
SP005
SP005_GCF_020616935.1
49c2
82c1
OG1RF
56c1 partial
93c4 partial
98c4 partial
121c1 partial
122c1 partial
136c1 partial
164c1 partial
179c1 partial
2.5 Kb
SP006
SP006_GCF_009662495.1
133c2 partial
173c1 partial
2.0 Kb
SP007
SP007_GCF_012594215.1
70c2
96c1
103c1
122c7
157c2
2.0 Kb
SP008
SP008_GCF_025958925.1
4c3 partial
33c1 partial
2.0 Kb
SP009
SP009_GCF_003030425.1
8c10
29c6
76c2
136c2
158c1
162c5
169c1
170c6
2.0 Kb
SP010
SP010_GCF_000211255.2
3c9 partial
2.0 Kb
SP011
SP011_GCF_025958805.1
144c1
2.0 Kb
epaR
EF2163 homolog
glycosyltransferase
glycosyltransferase
glycosyltransferase
glycosyltransferase
glycosyltransferase
glycosyltransferase
glycosyltransferase
glycosyltransferase
glycosyltransferase
glycosyltransferase
glycosyltransferase
Peptide O-xylosyltransferase
Wzy-like polymerase
Wzy-like polymerase
Wzx-like flippase
Wzx-like flippase
IspD/TarI-family cytidylyltransferase
TagD glycerol-3-phosphate cytidylyltransferase
LicD family protein
LicD family protein
LicD-like phosphotransferase
LicD-like protein
CDP-glycerol glycerophosphotransferase
CDP-glycerol glycerophosphotransferase
polysaccharide biosynthesis protein
polysaccharide biosynthesis protein
polysaccharide biosynthesis protein
O-antigen ligase family protein
O-antigen ligase family protein
O-antigen ligase family protein
O-antigen ligase family protein
epimerase/dehydratase
epimerase/dehydratase
DUF3329 domain-containing protein
DUF5776 domain-containing protein
DUF5776 domain-containing protein
DUF5776 domain-containing protein
DUF5776 domain-containing protein
DUF6056 domain-containing protein
DUF6056 domain-containing protein
DUF6056 domain-containing protein
GW-domain glycosaminoglycan-binding protein
GW-domain glycosaminoglycan-binding protein
GW-domain protein
GW-domain protein
GH25 family lysozyme
GH25 family lysozyme
glycosyl hydrolase
glycosyl hydrolase
lipoprotein
lipoprotein
membrane/transport protein
membrane/transport protein
transposase/mobile element
transposase/mobile element
hypothetical protein

### Figure S3

D

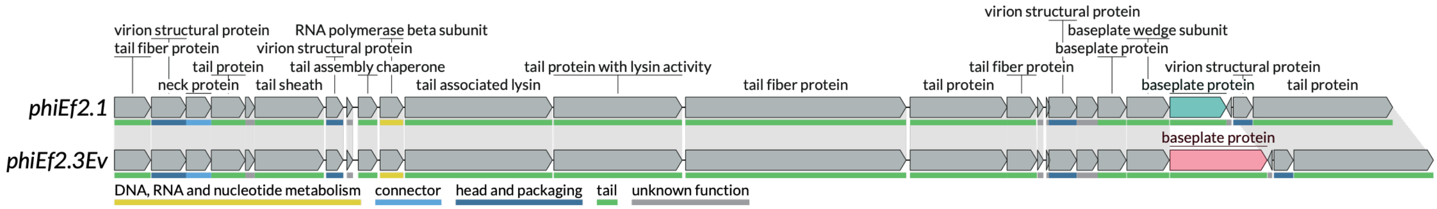


E

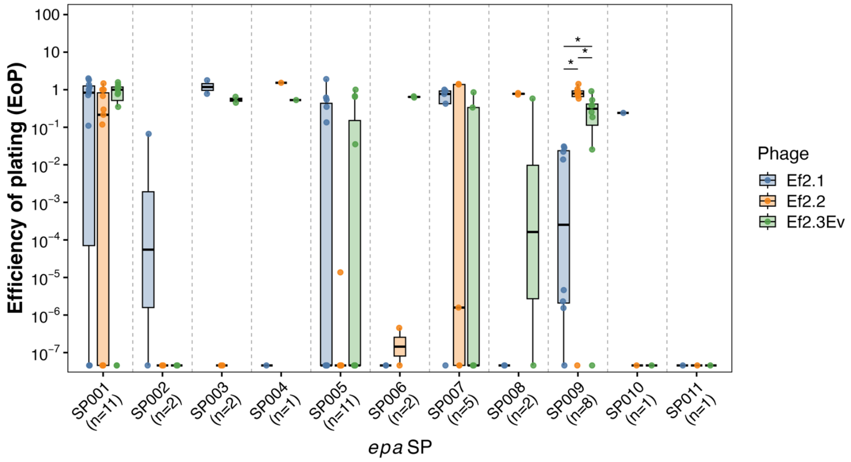


**Figure S3. Architecture-based classification of *epa* variable regions and associated phage phenotypes.**
(A) Neighbor-joining architecture-distance tree of the 367 quality-controlled *epaR*-to-EF2163 loci obtained from complete *E. faecalis* RefSeq genomes. Pairwise distances were calculated from ordered orthogroup strings using a weighted combination of orthogroup-content Jaccard distance and orthogroup-adjacency Jaccard distance. Colors indicate assignment to SP001–SP015. (B) Distribution of the 367 RefSeq genomes among the 15 *epa* variable-region SPs. Numbers above bars indicate the number of genomes assigned to each SP. (C) Locus-architecture comparisons between RefSeq SP representative loci and clinical *E. faecalis* isolates assigned to SP001–SP011. For each SP, the RefSeq representative locus is shown first; V583 is additionally shown for SP001. Clinical isolates with complete assignments are shown next, followed by partial or terminal assignments. CDSs are colored according to curated orthogroup annotations, and links connect CDSs assigned to the same curated orthogroup. (D) Comparative organization of the tail-associated genomic regions of ΦEf2.1 and ΦEf2.3Ev. Both phage genomes were re-annotated with Pharokka and Phold, and tail-associated regions were compared and visualized using LoVis4u. Predicted CDS functions are indicated above the loci, and functional gene modules are indicated below. The highlighted baseplate-associated CDSs represent a difference identified between the two phages within the otherwise largely conserved tail-associated regions. (E) EoP values for ΦEf2.1, ΦEf2.2, and ΦEf2.3Ev stratified by *epa* variable-region SP across all 46 strains assigned to an *epa* SP; two strains with undetermined assignments were excluded. Each point represents the mean EoP from three replicate assays for one strain. Statistical comparisons were restricted to SPs containing at least three strains. Within each eligible SP, overall differences among the three phages were evaluated using the Friedman test followed by paired Wilcoxon signed-rank tests for all three phage pairs. Pairwise *P* values within each SP were adjusted using the BH method. Asterisks indicate BH-adjusted pairwise *P* values: *, $P < 0.05$.

### Figure S4

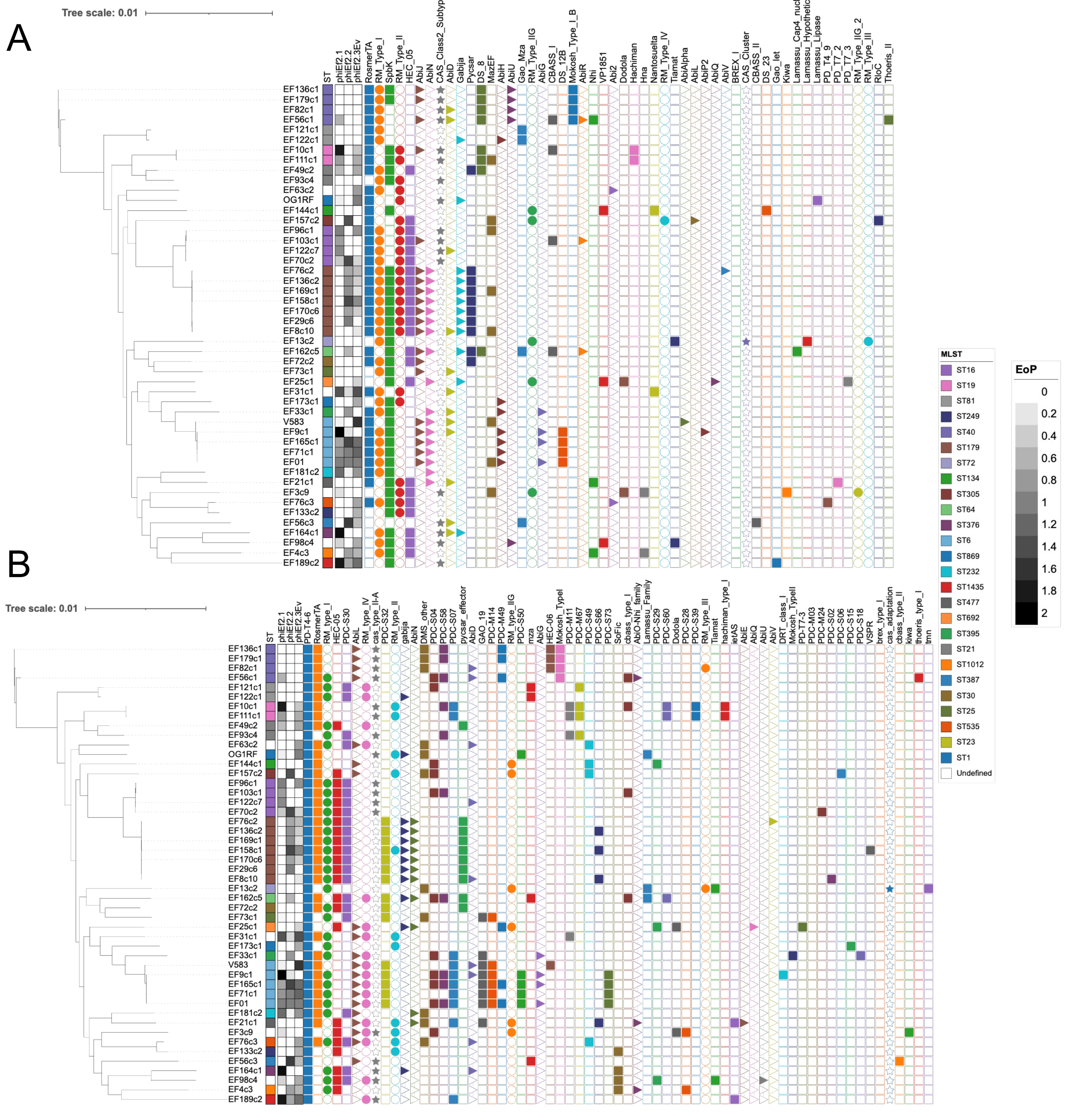


**Figure S4. Phylogenetic distribution of predicted anti-phage defense systems in the *Enterococcus faecalis* strain collection.**
Core-genome maximum-likelihood phylogeny of the 45 clinical isolates and three reference/control strains (EF01, OG1RF, and V583), annotated with multilocus sequence type, EoP values for ΦEf2.1, ΦEf2.2, and ΦEf2.3Ev, and predicted anti-phage defense-system profiles identified using (A) DefenseFinder and (B) PADLOC. Strains are ordered according to the core-genome phylogeny. EoP values are represented by the indicated grayscale. Colored symbols indicate detection of the corresponding predicted defense system, whereas empty symbols indicate absence. Defense-system nomenclature follows the respective DefenseFinder or PADLOC database.
